# Distinct lateral prefrontal regions are organized in an anterior-posterior functional gradient

**DOI:** 10.1101/2020.12.16.423034

**Authors:** Pin Kwang Tan, Cheng Tang, Roger Herikstad, Arunika Pillay, Camilo Libedinsky

**Affiliations:** The N1 Institute for Health, National University of Singapore (NUS), Singapore; Department of Psychology, Royal Holloway, University of London, Egham, United Kingdom; Department of Psychology, NUS, Singapore; Institute of Molecular and Cell Biology, A*STAR, Singapore

## Abstract

The dorsolateral prefrontal cortex (DLPFC) is composed of multiple anatomically-defined regions involved in higher-order cognitive processes, including working memory and selective attention. It is organized in an anterior-posterior global gradient, where posterior regions track changes in the environment, while anterior regions support abstract neural representations. However, whether the global gradient results from a smooth gradient that spans regions, or an overall trend emerging from the organized arrangement of functionally distinct regions is unknown. Here, we provide evidence to support the latter, by analyzing single-neuron activity along the DLPFC of non-human primates trained to perform a memory-guided saccade task with an interfering distractor. Additionally, we show that the posterior DLPFC plays a particularly important role in working memory, in sharp contrast with the lack of task-related responses in the anterior DLPFC. Our results validate the functional boundaries between anatomically-defined DLPFC regions and highlight the heterogeneity of functional properties across regions.

## Introduction

The realization that the cerebral cortex is parcellated into distinct interconnected brain regions is a cornerstone of our understanding of brain function. This idea was initially inspired by the observed segregation of cytoarchitectonic and functional properties across the cortical surface (Brodmann, 1909; Penfield and Jasper, 1954). Since then it has been further supported by multiple converging lines of evidence, including differences in anatomical and functional connectivity between regions (Glasser et al., 2016; Thomas Yeo et al., 2011; Yeterian et al., 2012). These anatomical and functional differences are two sides of the same coin, as differences in anatomy, such as inputs to the region, outputs to other regions, within-region connectivity, and intrinsic neuronal properties mould the functional properties of a specific brain region (Zylberberg and Strowbridge, 2017).

The dorsolateral prefrontal cortex (DLPFC, **Figure 1A**) has been parcellated into separate brain regions based on anatomical and functional properties (Kaping et al., 2011; Yeterian et al., 2012). It is involved in cognitive functions such as working memory, selective attention, and motor planning (Miller and Cohen, 2001). Substantial evidence supports the notion of a global functional gradient along the anterior-posterior axis of the DLPFC (Badre and D’Esposito, 2009, 2007; Koechlin et al., 2003; Petrides, 2005; Riley et al., 2018, 2017). This functional gradient appears to reflect a functional hierarchy, with anterior regions supporting more abstract neural representations and complex action rules, and posterior regions tracking moment-to-moment changes in the environment and the organism (Badre and D’Esposito, 2009; Constantinidis and Qi, 2018).

**Figure 1.**
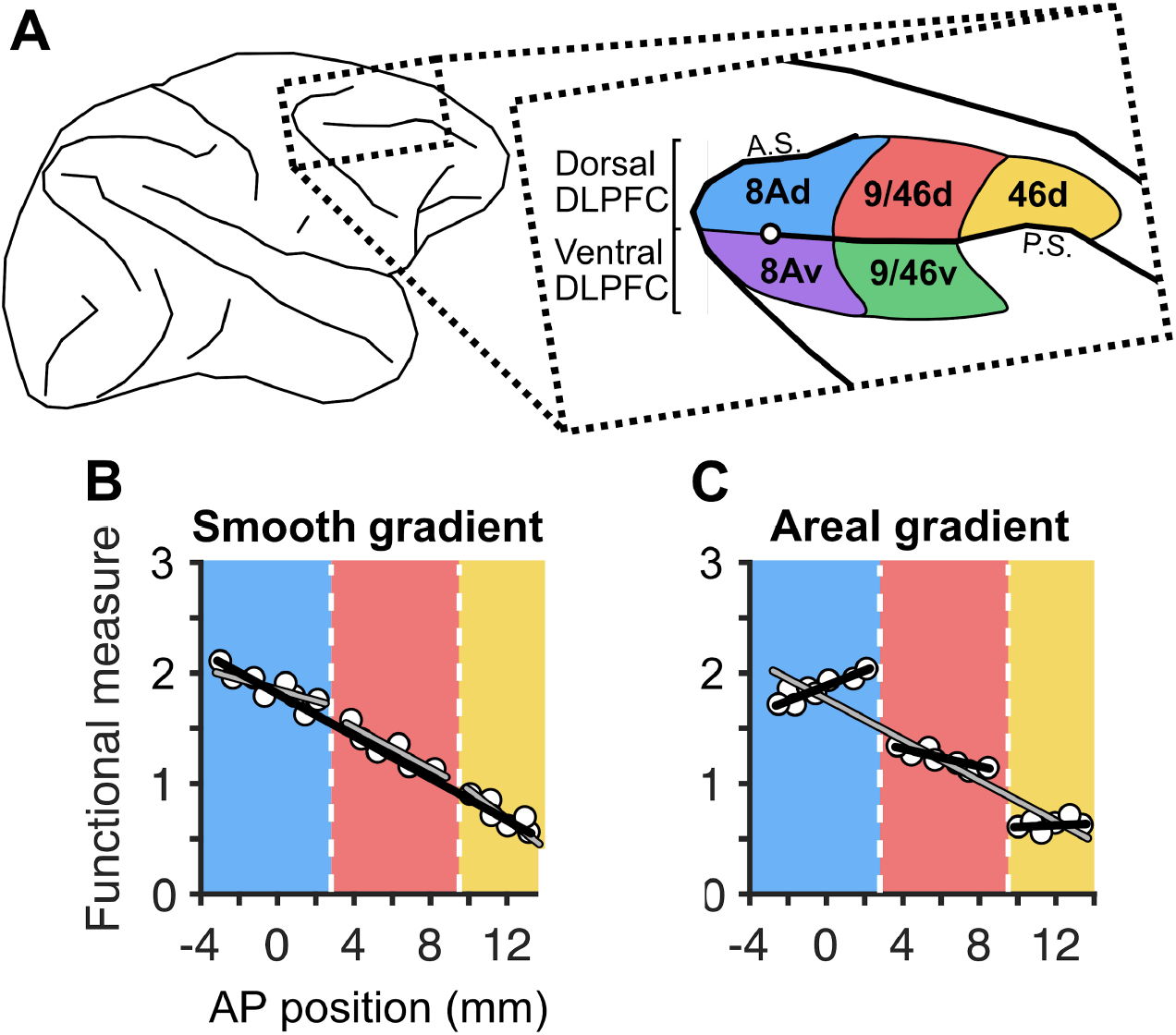
A global functional gradient can result from smooth or areal gradients along the anterior-posterior axis of the DLPFC. **A**, Schematic of the macaque brain, highlighting dorsal and ventral dorsolateral prefrontal regions (inset) with the posterior tip of the principal sulcus marked with a white circle (adapted from Petrides et al. 2012), A.S. and P.S. denote the arcuate sulcus and the principal sulcus., **B**, A smooth gradient of functional properties in the dorsal DLPFC, where a 1-segment linear model (black line) fits the data better than a 3-segment discontinuous piecewise model (grey lines), **C**, An areal gradient of functional properties in the dorsal DLPFC, where a 3-segment discontinuous piecewise model (black lines) fits the data better than a 1-segment linear model (gray line). Vertical dashed lines represent estimated anatomical boundaries between areas 8Ad, 9/46d, and 46d. Zero on the x-axis denotes the posterior tip of the principal sulcus.

Such global functional gradients in the DLPFC could be the result of a *smooth gradient* of functional properties along the anterior-posterior axis (**Figure 1B**). If this were true, then the segregation of DLPFC into different regions would be misleading, as a smooth gradient would instead suggest that the DLPFC acts as a single functional region, despite its heterogeneous anatomy. Alternatively, these global functional gradients could be the result of an overall trend that emerges from the organized arrangement of distinct areas, or an *areal gradient* (**Figure 1C**). If this were the case, then the segregation of DLPFC into different functional regions would be justified.

On the one hand, a smooth gradient could be expected based on the observation of what appears to be a smooth functional gradient along its anterior-posterior axis (Riley et al., 2018, 2017), as well as the high mixture of selectivities observed across DLPFC areas, which lack a clear topography (Mante et al., 2013; Rao et al., 1997; Rigotti et al., 2013). On the other hand, an areal gradient could be expected based on the observation of marked functional differences between DLPFC areas (Kaping et al., 2011), as well as the known anatomical differences between regions (Barbas, 2015; Markov et al., 2014; Petrides et al., 2012; Yeterian et al., 2012). Anatomically, the anterior-posterior axis of the DLPFC can be subdivided into three distinct areas - 8A, 9/46, and 46, from posterior to anterior - defined by specific cytoarchitectonic and connectivity patterns (Markov et al., 2014; Petrides et al., 2012; Yeterian et al., 2012). For example, area 9/46 receives input from the insula, while the adjacent area 46 does not (Yeterian et al., 2012). Furthermore, the principal sulcus appears to divide the regions into distinct dorsal and ventral portions with different connectivity patterns. For instance, the posterior cingulate cortex projects to the dorsal portion of these regions, while it does not project to their ventral portions (Yeterian et al., 2012). Such input differences may imply functional differences between these anatomically defined regions, and support an areal gradient across the DLPFC.

Determining whether the anterior-posterior functional global gradient in DLPFC is the result of a smooth gradient or an areal gradient will help constrain models of the prefrontal cortex, guide electrode positioning in future studies, as well as contextualize the conclusions of prior studies.

To assess whether the functional gradient across the DLPFC is the result of a smooth or an areal gradient, we recorded the activity of neurons in the DLPFC of two monkeys while they performed a working memory task that required ignoring a distractor shown mid-way through the delay period, i.e. between delays 1 and 2. We assessed the properties of single neurons as a function of their position along the anterior-posterior axis in individual monkeys. The physiological properties we explored included those during the visual target and distractor presentation periods (proportion of target and distractor selective cells, response latency, receptive field size, the strength of selectivity, and the degree of distractor filtering), and during both delay periods (last 500ms of a 1000ms delay; the proportion of target selective and nonlinear mixed selective cells during the delay, memory field sizes, and the strength of selectivity). We found that several of these measures had an anterior-posterior global gradient, and these were better explained by an areal gradient, rather than a smooth gradient. These results support the notion that global gradients are the result of the organized arrangement of distinct brain regions. To determine whether the different DLPFC regions perform different functions during the task, we analyzed the population level properties of the different functionally-defined areas. Specifically, we assessed regional information magnitude and stability across time of target and distractor location during correct and error trials. We observed that the posterior DLPFC plays a particularly important role in spatial working memory, in sharp contrast with the lack of task-related responses in the anterior DLPFC.

## Results

We recorded from the DLPFC of two adult macaque monkeys that were trained to perform a delayed saccade task with an intervening distractor (**Figure 2A**). The monkeys achieved a behavioral performance of ~75% correct trials (74% for Monkey A and 77% for Monkey B, across 4 recording sessions per monkey). Our analyses focused on physiological properties during the target and distractor presentation periods (300ms each; **Figure 2A**, red and green periods respectively), and the last 500ms of Delay 1 and Delay 2 periods (**Figure 2A**, yellow and orange periods respectively). We chose to analyze the last 500ms of the delay periods because we previously showed that the activity during the first ~500ms of the first delay had strong temporal dynamics, whereas the activity during the last 500ms was more stable (Parthasarathy et al., 2017).

**Figure 2.**
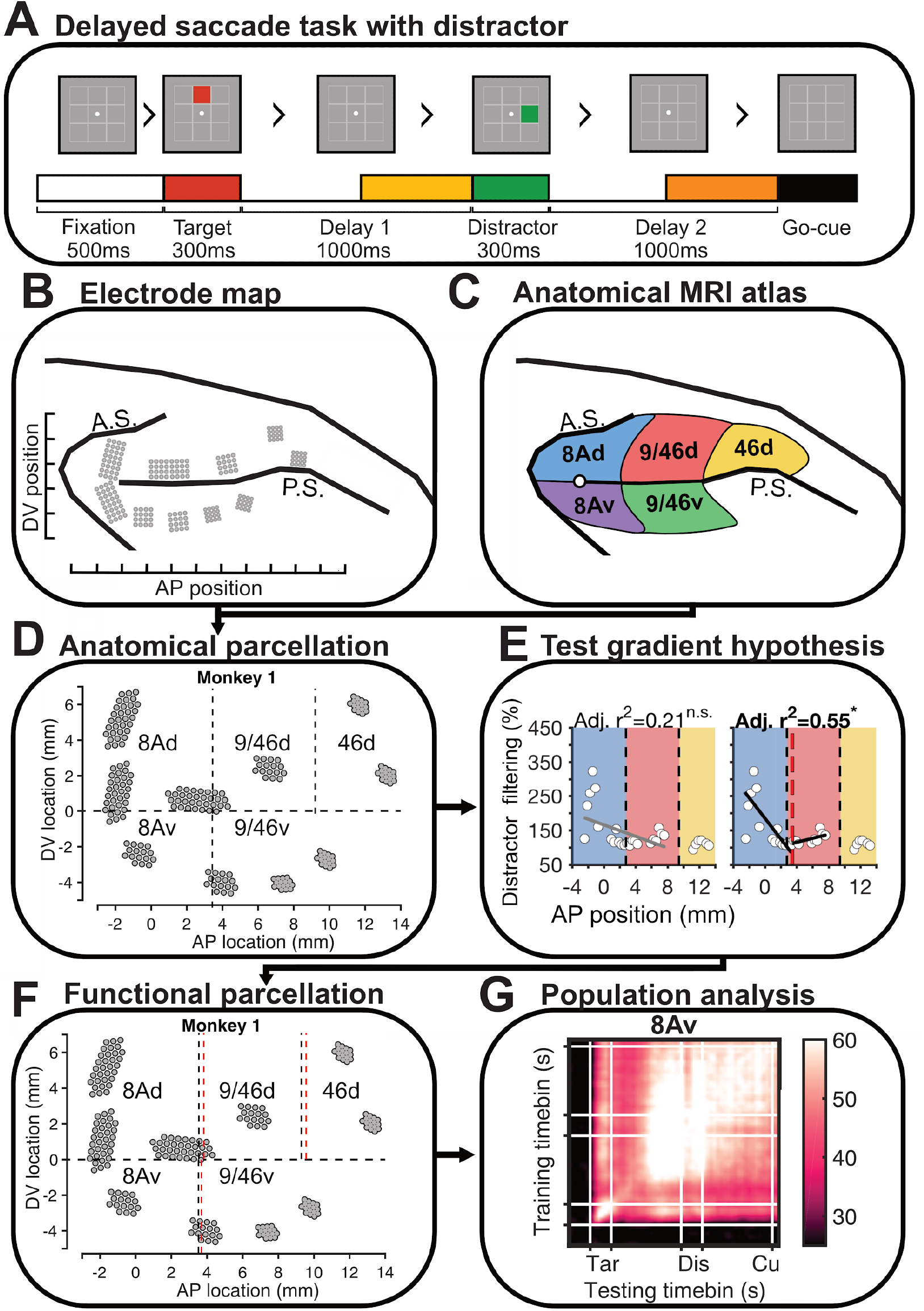
Overview of methods. **A**, The monkeys performed a delayed-saccade task with distractor interference. The periods analyzed are shown in the colored horizontal bars, including target (red), distractor (green), and delay (yellow and orange) periods; **B**, We recorded from the dorsolateral prefrontal cortex of 2 macaque monkeys *(Macaca fascicularis)* and estimated the anterior-posterior and dorso-ventral electrode positions from surgery images (not shown); **C**, We estimated native space anatomical parcellations using a neuroimaging atlas of *M. fascicularis;* **D**, Electrode positions in one monkey and anatomical parcellation boundaries (black dashed lines); **E**, To test whether the global gradient is the result of a smooth or an areal gradient across a range of functional measures, we tested whether a 1-segment or a 2-segment discontinuous piecewise model better described the data, separately for the dorsal and ventral DLPFC and separately for each monkey. At the top of each plot is shown the adjusted R^2^ of the 1- or 2-segment model fit (significant fits are highlighted with bold font and an asterisk). Regression lines are shown for the 1-segment model (left) and 2-segment model (right). The red vertical dashed line represents the estimated functional boundary between 8Ad and 9/46d for this measure, **F**, From the previous analysis, we derived functional parcellation boundaries (red dashed lines, black dashed lines are anatomical boundaries) and grouped neurons based on this functional parcellation map constructed for each monkey; **G**, Using the functional parcellations in F, we assessed population-level functional differences between regions using cross-temporal decoding analyses and measuring target information quantity (decoding performance) and stability across regions (neurons pooled across both monkeys).

To examine gradients along the anterior-posterior axis, we first estimated the anterior-posterior position (AP position) of each microelectrode (and hence neuron) from images taken during surgery (**Figure 2B**). The AP position of each neuron was calculated with reference to the posterior tip of the principal sulcus, where zero denotes the posterior tip of the principal sulcus, negative values represent locations posterior to the tip, and positive values represent locations anterior to the tip. We grouped electrodes into either the dorsal or ventral DLPFC (dDLPFC and vDLPFC respectively) using the principal sulcus as the dividing line. We estimated the anatomical boundaries of each region (dorsal and ventral areas 8A, 9/46, 46) from a macaque MRI atlas (Frey et al., 2011)(**Figure 2C**). Combining the AP positions of electrodes and the anatomical boundaries, we parcellated electrodes into dDLPFC and vDLPFC subregions, separately for each monkey (**Figure 2D**). We recorded a total of 320 neurons from the dDLPFC (comprising areas 8Ad, 9/46d, and 46d) and 169 from the vDLPFC (comprising areas 8Av and 9/46v). Areas 9/46v and 46d contain data from only one monkey (Monkey 1).

### 3.1 Functional organization of single neurons along the anterior-posterior axis

First, we sought to test whether the functional properties of DLPFC neurons were organized as a global gradient along the anterior-posterior axis. For the dDLPFC and vDLPFC, we obtained physiological measures of single neurons and plotted them in the y-axis, and their anterior-posterior anatomical position was plotted in the x-axis, regardless of their dorso-ventral position within the region (**Figure 2E**). The data was spatially smoothed using a 0.5mm sliding window with 0.17mm overlap between windows (number of neurons per point for Monkey 1 = 1-56, mean = 11.9; Monkey 2 = 1-11, mean = 5.7; histogram of neuron counts per point in **Figure 2 - figure supplement 1**).

To assess whether there was a significant global gradient along the anterior-posterior axis we fitted a 1-segment linear model to the data (henceforth referred to as a 1-segment model), separately for the dorsal and ventral DLPFC (**Figure 2E, left;** see full statistical flowchart in **Figure 2 - figure supplement 2**). We then compared its adjusted R^2^ to a distribution of R^2^s obtained from models fit on data shuffled along the anterior-posterior axis (1000 iterations). 1-segment models with adjusted R^2^ values that exceeded the 95^th^ percentile of the shuffled distribution were considered to have significant global gradients. In the dDLPFC, out of the 13 functional measures assessed, 12 of them showed a global gradient in one or both monkeys (all measures except for the proportion of neurons with non-linear mixed selectivity) (**Figure 3C-H and 4C-I, columns i and iii**). In contrast, in the vDLPFC, out of the 13 functional measures assessed, only 2 of them (proportion of neurons with non-linear mixed selectivity, and selectivity index in Delay 1) showed a global gradient in monkey 1 (this could not be assessed in monkey 2, since we did not record from area 9/46v) (**Figures 5C-H and 6E, H - column i**).

**Figure 3.**
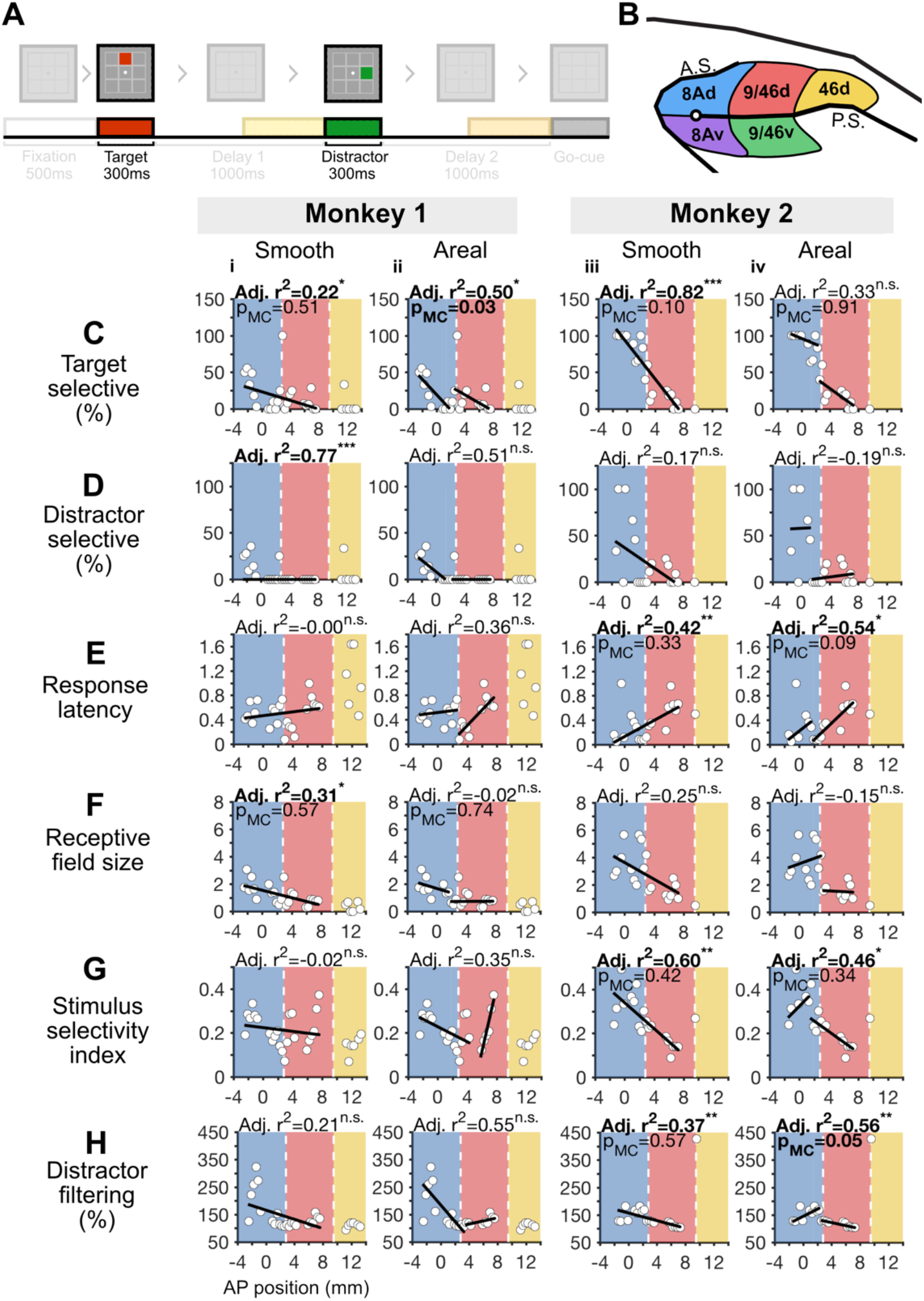
Analysis of anterior-posterior gradients in the dDLPFC during the stimuli presentation periods. **A**, Task design highlighting the target (red bar) and distractor (green bar) periods used in this figure. **B**, Dorsolateral prefrontal cortex, with color-coded regions that match the color-coded figures below in C-H (blue, red, and yellow denoting areas 8Ad, 9/46d, and 46d respectively)**. C-H**, We fit 1 - and 2-segment models spanning areas 8Ad and 9/46d (area 46d because was not included here because we did not record from this region in Monkey 2, see **Figure 3 - figure supplement 1** for plots that include area 46d). From left to right, the columns of plots depict the 1-segment model for Monkey 1, the 2-segment model for Monkey 1, the 1-segment model for Monkey 2, and the 2-segment model for Monkey 2. Model fits that are significant versus a shuffled baseline are denoted with an adjusted R^2^ in bold and asterisks (n.s. = not significant). For each monkey, we compared the model fits of the 1- and 2-segment models versus the alternative model. For instance, to compare a 1-segment model (Adj. R^2^=0.22) against the 2-segment model (Adj. R^2^=0.50), we calculated the probability of obtaining a 1-segment model fit with an adj. R^2^ of 0.22, when fit on a surrogate dataset with a 2-segment adjusted adj. R^2^ of 0.50, and with the same mean and standard deviation as the original data. The significance value of the model-comparison is shown in the plot. Note that the model comparison step was not conducted for distractor selectivity % for monkey 1, as it was not possible to generate surrogate datasets that met the desired criterion.

**Figure 4.**
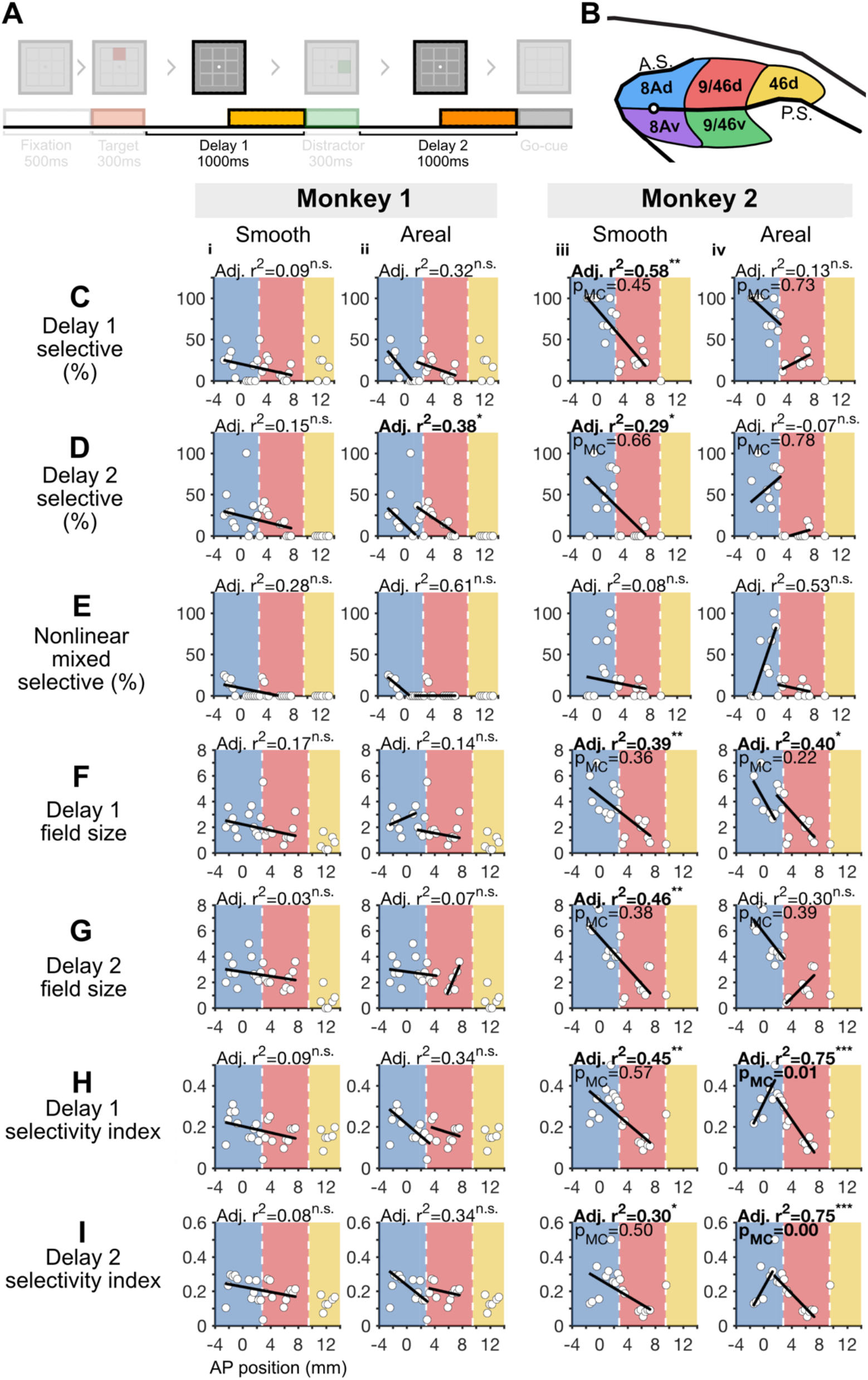
Analysis of anterior-posterior gradients in the dDLPFC during the delay periods. **A**, Task design, with the analyzed period highlighted (delay periods). **B**, Dorsolateral prefrontal cortex, with color-coded regions that match the color-coded figures below in C, i.e. blue, red and yellow denoting 8Ad, 9/46d, and 46d respectively. **C-I**, We fit 1- and 2-segment gradients across the anterior-posterior axis of the dorsal DLPFC for delay period measures (area 46d because was not included here because we did not record from this region in Monkey 2, see **Figure 4 - figure supplement 1** for plots that include area 46d). For a description of the plots see the legend of **Figure 3C**.

**Figure 5.**
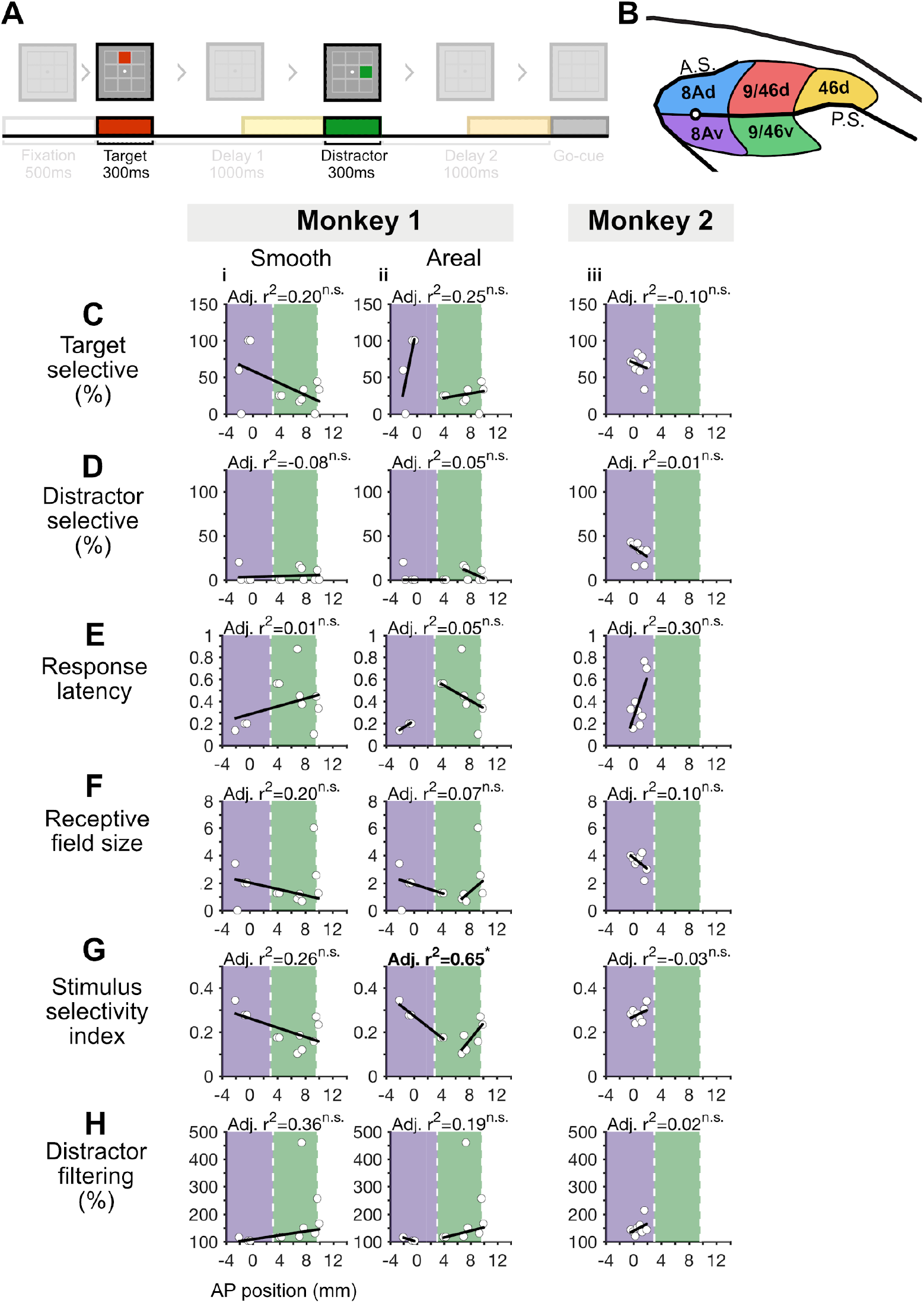
Analysis of anterior-posterior gradients in vDLPFC during the stimuli presentation periods. **A**, Task design, with the analyzed period highlighted (target period). **B**, Dorsolateral prefrontal cortex, with color-coded regions that match the color-coded figures below in C, i.e. purple and green denoting 8Av and 9/46v respectively. **C**, We fit 1- and 2-segment gradients across the anterior-posterior axis of the ventral DLPFC for target period measures, with the two leftmost columns (displaying the 1-segment and 2-segment model fits) belonging to monkey 1 and the two rightmost to monkey 2. For a description of the plots see the legend of **Figure 3C**.

**Figure 6.**
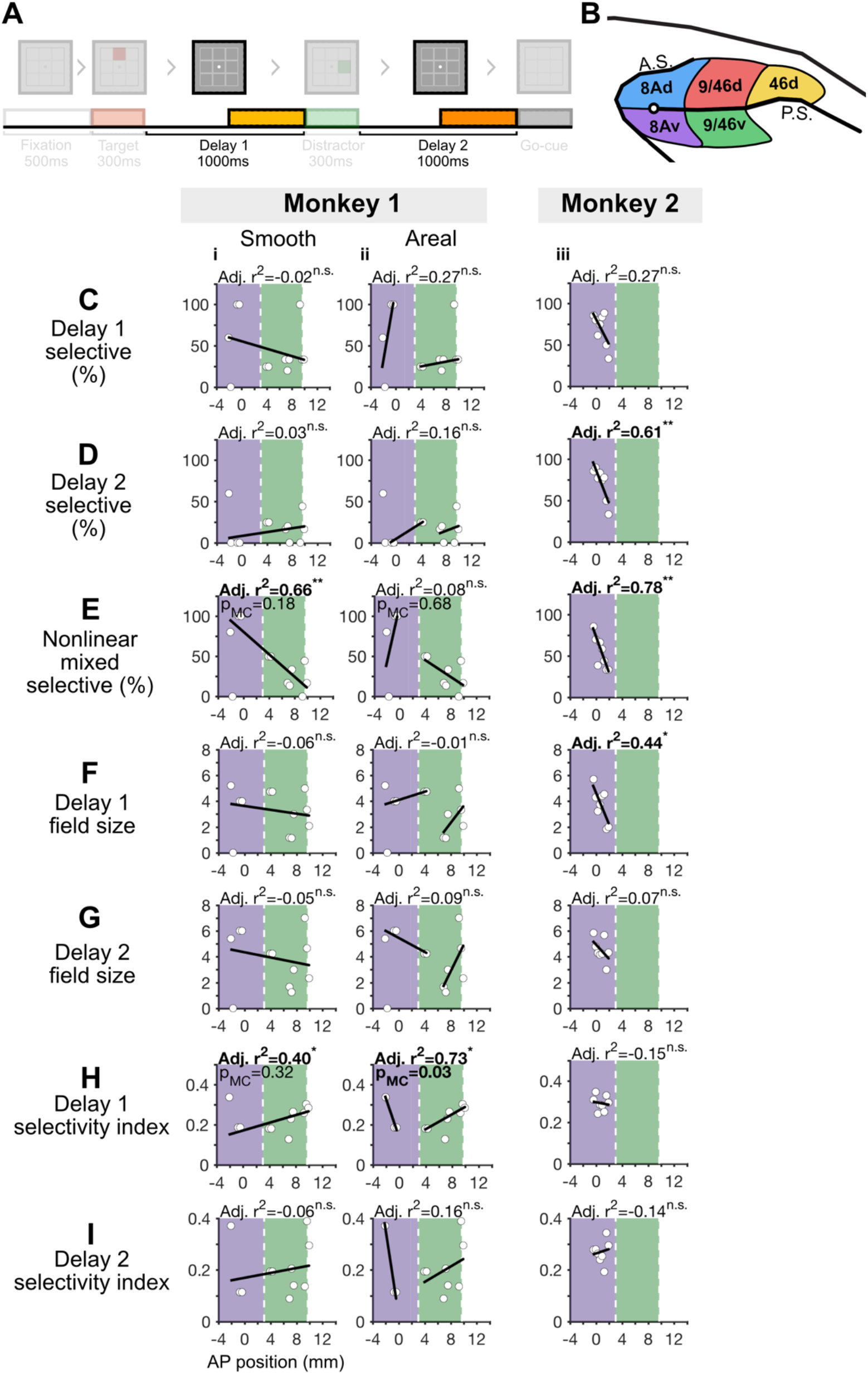
Analysis of anterior-posterior gradients in vDLPFC during the delay periods. **A**, Task design, with the analyzed period highlighted (delay period). **B**, Dorsolateral prefrontal cortex, with color-coded regions that match the color-coded figures below in C, i.e. purple and green denoting 8Av and 9/46v respectively. **C**, We fit 1- and 2-segment gradients across the anterior-posterior axis of the ventral DLPFC for delay period measures, with the two leftmost columns (displaying the 1-segment and 2-segment model fits) belonging to monkey 1 and the two rightmost to monkey 2. For a description of the plots see the legend of **Figure 3C-I**.

Then, we aimed to determine whether those functional measures that were organized as a global gradient were better explained by a smooth or an areal gradient. To this end, we carried out the following analysis.

First, we fitted a 2-segment discontinuous piecewise linear model to the data (henceforth referred to as 2-segment model) (Ryan and Porth, 2007), where each ‘break’ corresponded to an estimated functional boundary (**Figure 2E, right**). The estimated functional boundaries were constrained by anatomical parcellations extracted from a macaque MRI atlas (Frey et al., 2011). Specifically, we allowed functional boundaries to deviate ±1.5mm from these anatomical boundaries to allow for inter-animal variations (Xu et al., 2018). This was done separately for the dDLPFC and vDLPFC, and for each monkey (**Figure 2F**). The goodness of fit of the 2-segment models was measured using the adjusted R^2^, which contains a penalty term for the increased number of parameters used in the piecewise models (Ryan and Porth, 2007). To assess the significance of a 2-segment model’s fit in the same manner as above, we compared its adjusted R^2^ to a distribution of adjusted R^2^s obtained from models fit on data shuffled along the anterior-posterior axis (1000 iterations). Models with adjusted R^2^ values that exceeded the 95^th^ percentile of the shuffled distribution were considered significant. In the dDLPFC, out of the 11 functional measures that showed a global gradient, 7 of them showed a significant 2-segment fit (**Figure 3C-H and 4C-I, columns ii and iv**). On the other hand, in the vDLPFC only one of the two functional measures that showed a global gradient, the Delay 1 selectivity Index, showed a significant 2-segment model adjusted R^2^ (**Figures 6H**).

Second, for each adjusted R^2^ value obtained (both 1- and 2-segment), we calculated the probability of obtaining that value or higher, given the adjusted R^2^ of the alternative model. For example, in Monkey 1, the R^2^ of the 1-segment model fitted to the proportion of selective cells during the target presentation period was 0.22, which was significantly higher than chance (**Figure 3Ci**). The fit obtained with a 2-segment model was 0.50, which was also significantly higher than chance (**Figure 3Cii**). Given this data, we asked what was the probability of obtaining a 1-segment model fit with an R^2^ of 0.22 or higher, given a *2-segment surrogate dataset* with a similar adjusted R^2^ as the 2-segment adjusted R^2^ of 0.50, and with a similar mean and standard deviation as the original data. This process was repeated for 1000 of such 2-segment surrogate datasets to create a distribution. The same procedure was followed for the 2-segment model fit with a *1-segment surrogate dataset* (with a similar R^2^ as the 1-segment fit, and a similar mean and standard deviation on the original data). For each model fit, we then obtained a *model-comparison p-value* (*p*_MC_) by comparing the actual adjusted R^2^ with the adjusted R^2^ distribution (see full statistical flowchart in **Figure 2 - figure supplement 2**). These model-comparison *p*-values are shown in **Figures 3-6** below the adjusted R^2^ values.

Third, we hypothesized that if the 1- or 2-segment model explains the data better than the other, its model-comparison *p*-value distribution would not be Gaussian (with a mean of 0.5), but rather it would have a distribution with a peak frequency biased towards zero (i.e. Weibull distribution biased towards 0). Thus, we used the Anderson-Darling test to assess whether either the 1-segment or 2-segment distribution followed a Weibull distribution with a bias towards 0. We included only measures with a significant global gradient, i.e. significant 1-segment fit versus noise. As described in more detail below, we found that the 2-segment model, but not the 1-segment model, is consistent with a Weibull distribution biased towards 0. Thus, the evidence supports the view that the global gradients observed are the result of areal gradients of functional properties (**Figure 7**).

**Figure 7.**
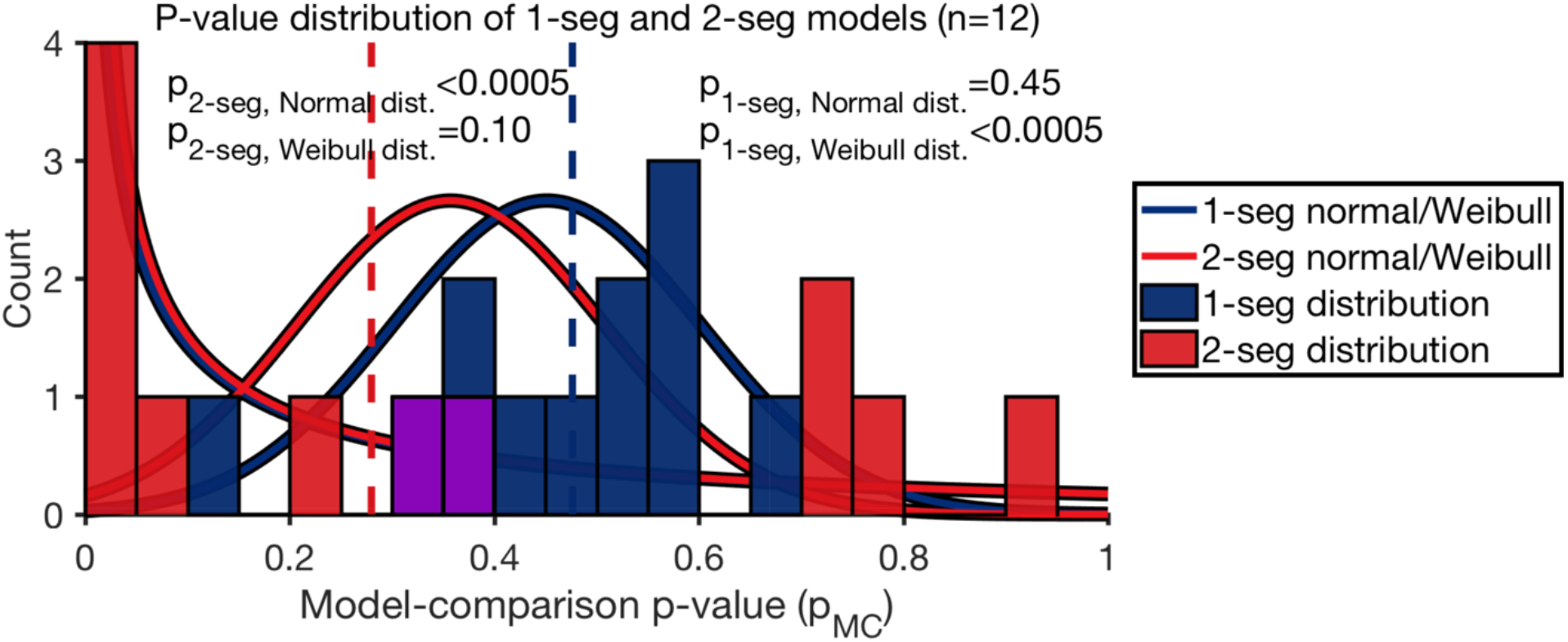
An areal gradient better describes the global gradient across the dDLPFC. Data distributions of model-comparison *p*-values for the 1-segment (blue bars) and 2-segment (red bars) models (overlap at 0.3 and 0.4 shown as purple bars), medians shown in the dashed lines in the respective colors. The Anderson-Darling test (AD-test) *p*-values are shown for testing whether the distributions can be described by a normal (μ_1-seg_=0.45, μ_2-seg_=0. 36, σ=0.15) or a Weibull (k=0.50, λ=0.50) distribution (colored according to the colors of the 1-segment and 2-segment data distributions). Note that the Weibull distribution models for the 1-segment and 2-segment distributions are identical but are slightly displaced horizontally for visualization purposes. Medians are denoted with a dashed line.

#### 3.1.1 Global Gradients in the dDLPFC

The following sections describe the global gradients of functional properties along the anterior-posterior axis during the stimuli and delay periods (**Figures 3-4**). Based on estimated anatomical parcellations, our electrodes spanned 3 regions of the dorsal DLPFC: 8Ad, 9/46d, and 46d (**Figure 3B**).

##### 3.1.1.1 Responses during stimuli presentation

First, we analyzed response properties during the presentation of the stimuli (target and distractor), namely the proportion of cells selective to target or distractor locations, response latency, receptive field size, stimulus selectivity, and degree of distractor filtering (**Figure 3C-H**).

*The proportion of target selective cells* was defined as the proportion of cells with a significant selectivity to the target location, during target presentation (ANOVA *p*-value < 0.05). In both monkeys, we found evidence for global gradients, such that the proportion of target selective cells decreased in more anterior regions (**Figure 3Ci, iii**). In Monkey 1 the global gradient was better explained by an areal gradient (*p*_Mc 1-seg_=0.51, *p*_MC 2-seg_ = 0.03, **Figure 3Ci, ii**). Including area 46d in the analysis revealed a significant global gradient, such that the proportion of target selective cells decreased in more anterior regions (**Figure 3 - figure supplement 1Ai**). This global gradient was better explained by an areal gradient (*p*_MC 1-v-2 seg_ = 0.24, *p*_MC 1-v-3 seg_ = 0.16, *p*_MC 2-v-1 seg_ < 0.01, *p*_MC 3-v-1 seg_ = 0.01, **Figure 3 - figure supplement 1Ai-iii**). In Monkey 2 it was inconclusive whether it was better explained by a smooth or an areal gradient (*p*_MC 1-seg_ = 0.10, *p*_MC 2-seg_ = 0.91, **Figure 3Ciii, iv**).

*The proportion of distractor selective cells* was defined as the proportion of cells with a significant selectivity to distractor location, during distractor presentation (ANOVA *p*-value < 0.05). In Monkey 1 we found evidence for a global gradient, such that the proportion of distractor selective cells decreased in more anterior regions (**Figure 3Di**). However, it was inconclusive whether the global gradient was better explained by a smooth or an areal gradient as we were unable to generate 1-segment surrogate datasets that met our criterion (**Figure 3Di, ii**). In Monkey 2 we found no evidence for a global gradient (**Figure 3Diii**).

*Response latency* was defined as the earliest responsive time-bin for each neuron’s preferred target location compared to baseline (−500ms to 0ms prior to target presentation, t-test *p*-value < 0.05). In Monkey 1 we found no evidence for a global gradient (**Figure 3Ei**). In Monkey 2 we also found evidence for a global gradient with longer latencies in more anterior regions but it was inconclusive whether it was better explained by a smooth or areal gradient (*p*_MC 1-seg_ = 0.33, *p*_MC 2-seg_ = 0.09, **Figure 3Eiii, iv**).

The *receptive field size* was defined as the number of locations where activity during target presentation was higher than the baseline (−500ms to 0ms prior to target presentation, t-test *p*-value < 0.05). In Monkey 1, we found evidence for global gradients, such that receptive field sizes decreased in more anterior regions (**Figure 3Fi**). However, it was inconclusive whether the global gradient was better explained by a smooth or an areal gradient (*p*_MC 1-seg_ = 0.57, *p*_MC 2-seg_ = 0.74, **Figure 3Fi, ii**). Including area 46d in the analysis revealed a significant global gradient, such that the receptive field sizes decreased in more anterior regions (**Figure 3 - figure supplement 1Di**). Despite this, it was inconclusive whether the global gradient was better explained by a smooth or an areal gradient (*p*_MC 1-v-2 seg_ = 0.91, *p*_MC 1-v-3 seg_ = 1.00, *p*_MC 2-v-1 seg_ = 0.78, *p*_MC 3-v-1 seg_ = 0.79, **Figure 3 - figure supplement 1Di-iii**). In Monkey 2 we found no evidence for a global gradient (**Figure 3Fiii**).

The *stimulus selectivity index* was defined as (*max*-*min*)/(*max*+*min*), where *max* is the average firing rate for the target location with the highest firing rate and *min* is the average firing rate for the target location with the lowest firing rate. In Monkey 1 we found no evidence of a global gradient (**Figure 3Gi**). In Monkey 2 we found evidence for a global gradient, with decreasing stimulus selectivity indices in more anterior regions, but it was inconclusive whether it was better explained by a smooth or an areal gradient (*p*_MC 1-seg_ = 0.42, *p*_MC 2-seg_ = 0.34, **Figure 3Giii, iv**).

*Distractor filtering* was defined as the ratio of mean activity evoked by the preferred target (maximum average firing rate) over the mean activity for a distractor in the same location. A value of 100 means that the target and distractor evoked the same activity, while a value higher than 100 means that the target evoked a larger response than the distractor (i.e., the distractor was filtered). In Monkey 1 we found no evidence for a global gradient (**Figure 3Hi**). In Monkey 2 we found evidence for a global gradient, with decreased filtering in more anterior regions, and this global gradient was better explained by a smooth or areal gradient (*p*_MC 1-seg_=0.57, *p*_MC 2-seg_ = 4.5 × 10^-2^, **Figure 3Hiii, iv**).

##### 3.1.1.2 Responses during the delay periods

Next, we analyzed response properties during the delay periods, namely the proportion of selective cells, memory field sizes, and target selectivity (**Figure 4**).

*The proportion of Delay 1 selective cells* was defined as the proportion of cells with a significant selectivity to the target location, during the last 500ms of the Delay 1 period (ANOVA *p*-value < 0.05). In Monkey 1 we found no evidence of a global gradient (**Figure 4Ci**). In Monkey 2 we found evidence for a global gradient, with a decreasing proportion of Delay 1 selective cells in more anterior regions (**Figure 4Ciii**). However, it was inconclusive whether the global gradient was better explained by a smooth or an areal gradient (*p*_MC 1-seg_ = 0.45, *p*_MC 2-seg_ = 0.73, **Figure 4Ciii, iv**).

The *proportion of Delay 2 selective cells* was defined as the proportion of cells with a significant selectivity to the target location, during the last 500ms of the Delay 2 period (ANOVA *p*-value < 0.05). In Monkey 1 we found no evidence for a global gradient (**Figure 4Di**). Including area 46d in the analysis revealed a significant global gradient, such that the proportion of Delay 2 selective cells decreased in more anterior regions (**Figure 4 - figure supplement 1Bi**). However, it was inconclusive whether the global gradient was better explained by a smooth or an areal gradient (*p*_MC 1-v-2 seg_ = 0.34, *p*_MC 1-v-3 seg_ = 0.48, *p*_MC 2-v-1 seg_ = 0.10, *p*_MC 3-v-1 seg_ = 0.08, **Figure 4 - figure supplement 1Bi-iii**). In Monkey 2 we found a global gradient with a decreasing proportion of Delay 2 selective cells in more anterior regions (**Figure 4Diii**). However, it was inconclusive whether the global gradient was better explained by a smooth or an areal gradient (*p*_MC 1-seg_ = 0.66, *p*_MC 2-seg_ = 0.78, **Figure 4Diii, iv**).

*The proportion of nonlinear mixed-selective cells* was defined as the proportion of cells with a significant interaction between target location and delay period (2-way ANOVA; factor-1: target location, factor-2: Delay 1 or Delay 2; interaction *p*-value < 0.05) (Parthasarathy et al., 2017). We found no evidence of global gradients in either monkey (**Figure 4Ei, iii**).

*Delay 1 field size* was defined as the number of locations where activity during the last 500ms of the Delay 1 period was higher than the baseline (−500ms to 0ms prior to target presentation, t-test *p*-value < 0.05). In Monkey 1 we found no evidence for a global gradient (**Figure 4Fi**). In Monkey 2, we found evidence for global gradients, with smaller Delay 1 field sizes in more anterior regions (**Figure 4Fiii**). However, it was inconclusive whether the global gradient was better explained by a smooth or an areal gradient (*p*_MC 1-seg_ = 0.36, *p*_MC 2-seg_ = 0.22, **Figure 4Fiii, iv**).

*Delay 2 field size* was defined as the number of locations where activity during the last 500ms of the Delay 2 period was higher than the baseline (−500ms to 0ms prior to target presentation, t-test *p*-value < 0.05). In Monkey 1 we found no evidence of a global gradient (**Figure 4Gi**). In Monkey 2 we found similar evidence for a global gradient, such that delay 2 field sizes decreased in more anterior regions (**Figure 4Giii**). But it was inconclusive whether the global gradient was better explained by a smooth or an areal gradient (*p*_MC 1-seg_ = 0.38, *p*_MC 2-seg_ = 0.39, **Figure 4Giii, iv**).

*Delay 1 selectivity index* was defined as *(max-min)/(max+min)*, where *max* is the average firing rate for the target location with the highest firing rate during the last 500ms of Delay 1 period, and *min* is the average firing rate for the target location with the lowest firing rate. In Monkey 1 we found no evidence for a global gradient (**Figure 4Hi**). In Monkey 2 we found evidence for a global gradient, such that delay 1 selectivity index decreased in more anterior regions (**Figure 4Hiii**). This global gradient was better explained by an areal gradient (*p*_MC 1-seg_=0.57, *p*_MC 2-seg_=0.01, **Figure 4Hiii**).

*Delay 2 selectivity index* was defined in the same manner as Delay 1 selectivity index, but for the last 500ms of Delay 2 period. In Monkey 1 we found no evidence of a global gradient (**Figure 4Ii**). In Monkey 2 we also found evidence for a global gradient, such that Delay 2 selectivity index became smaller in more anterior regions (**Figure 4Iiii**). This global gradient was better explained by an areal gradient (*p*_MC 1-seg_=0.50, *p*_MC 2-seg_<0.01, **Figure 4Iiii**).

In summary, dorsal DLPFC regions consistently show evidence of global functional gradients along the anterior-posterior axis, such that anterior regions had less target information in both target and delay periods. Importantly, the direction of the gradients in both monkeys was consistent for all the functional measures assessed, lending strong support for the existence of these global gradients.

#### 3.1.2 Global Gradients in the vDLPFC

Based on the estimated anatomical parcellation of the vDLPFC, our electrodes spanned 2 regions in Monkey 1 (8Av and 9/46v) and only 1 region in Monkey 2 (8Av)(**Figure 1A**). The following sections describe the stimulus and delay period functional properties of these regions. Note that only Monkey 1’s data can be tested for global gradients because Monkey 2 has electrodes in area 8Av only. Nevertheless, the model fits for Monkey 2 are included for visual comparison.

##### 3.1.2.1 Responses during the stimulus period

We characterized the same functional properties during the target and distractor period in the vDLPFC as those we used to characterize the dDLPFC (**Figure 5**).

Unlike in the dorsal DLPFC, where Monkey 1 showed global gradients in 3 out of the 7 functional measures, and Monkey 2 showed global gradients in 5 out of the 7 functional measures, in the ventral DLPFC of Monkey 1 we found no evidence of a global gradient in any of the functional measures assessed (**Figure 5G**).

##### 3.1.2.2 Responses during the delay period

We characterized the same functional properties during the delay periods in the vDLPFC as those we used to characterize the dDLPFC (**Figure 6**).

We found global gradients in Monkey 1 in the proportion of nonlinear mixed selective cells, and the Delay 1 selectivity index (**Figure 6E, H**). The proportion of nonlinear mixed-selective cells decreased in more anterior regions (**Figure 6Ei**). But it was inconclusive whether the global gradient was better explained by a smooth or an areal gradient in Monkey 1 (*p*_MC 1-seg_=0.18, *p*_MC 2-seg_=0.68, **Figure 6Ei, ii**). The Delay 1 selectivity index increased in more anterior regions (**Figure 6H**), and this global gradient was better explained by an areal gradient (*p*_MC 1-seg_=0.32, *p*_MC 2-seg_=0.03, **Figure 6Hii**).

In summary, evidence for the existence of global gradients in the ventral DLPFC regions is less reliable than in the dorsal regions. However, the data originates from a single monkey, so we are unable to make a strong claim about the presence or absence of gradients in these ventral regions.

#### 3.1.2 Global gradients are better explained by areal gradients

The previous analyses revealed that a number of functional measures were better explained by a 2-segment model than a 1-segment model (*p*_MC 2-seg_ < 0.05, and *p*_MC 1-seg_ > 0.05). Here we asked whether this trend was true across measures, by analyzing the *p*_MC_ distribution across all functional measures with a significant global gradient (**Figure 7**). If the 1-segment model describes the data well across the functional measures, then the *p*_MC 1-seg_ distribution should be biased towards 0, whereas if it does not describe the data well, the *p*_MC 1-seg_ distribution should follow a normal distribution. Similarly for the 2-segment model.

To determine whether the 1- or 2-segment model describes the data better, we used the Anderson-Darling test (AD-test) to determine whether their *p*_MC_ distributions follow a Weibull distribution with a bias towards 0, or a normal distribution (**Figure 7 - figure supplement 1)**. For the AD-test a *p*-value > 0.05 means that we cannot reject the hypothesis that the data has the tested distribution, while a p-value < 0.05 means that we can reject that hypothesis. We carried out all tests using 3 sets of parameters for each Weibull/normal distribution to allow for flexibility in fitting to the *p*_MC_ distribution (λ=0.25, 0.50 or 0.75; σ=0.10, 0.15 or 0.30; λ=Weibull scale parameter, σ=normal standard deviation parameter).

We found that neither the 1-segment nor the 2-segment data had a normal distribution with μ=0.50 (σ=0.10, 0.15, 0.30; AD test, all *p*<0.05). Hence we tested if normal distributions centered on the respective means of the data described either distribution (μ fixed at each data distribution’s mean, μ_1-seg_=0.36, μ_2-seg_= 0.45, μ=normal mean parameter, see **Figure 7 - figure supplement 1**). Similarly, we tested if Weibull distributions biased towards 0 described either distribution (k fixed at k=0.50, k=Weibull shape parameter, see **Figure 7 - figure supplement 1**).

We found that we cannot reject the hypothesis that the 1-segment model has a normal distribution (μ=0.45, σ=0.15, *p*_1seg, normal_=0.45, AD-stat_1seg, normal_ =0.34, **Figure 7** *blue distribution*), whereas we can reject the hypothesis that the 2-segment model has a normal distribution (μ=0.36, σ=0.15, *p*_2seg, normal_<5 × 10^-4^, AD-stat_2seg, normal_=8.11, **Figure 7** *red distribution*). On the other hand, we can reject the hypothesis that the 1-segment model has a Weibull distribution biased towards zero (k=0.50, λ=0.50, *p*_1seg, weibull_<5 × 10^−4^, AD-stat_1seg, Weibull_=3.54, **Figure 7** *blue distribution)*, whereas we cannot reject the hypothesis that the 2-segment model has a Weibull distribution biased towards zero (k=0.50, λ=0.50, *p*_2seg, Weibull_=0.10, AD-stat_2seg, Weibull_=0.62, **Figure 7** *red distribution).* Variations of the σ and λ parameters led to rejections of all the hypotheses (**1-segment, normal**: μ=0.45, σ=0.10, *p*_1seg, normal_=0.02, AD-stat_1seg, normal_=0.87; μ=0.45, σ=0.30, *p*_1seg, normal_=1.2 × 10^-3^, AD-stat_1seg, normal_=1.29; **1-segment, Weibull**: k=0.50, λ=0.25, *p*_1seg, Weibull_<5 × 10^-4^ , AD-stat_1seg, Weibull_=4.39; k=0.50, λ=0.75, *p*_1seg, Weibull_<5 × 10^-4^ , AD-stat_1seg, Weibull_=2.87; **2-segment, normal**: μ=0.36, σ=0.10, *p*_2seg, normal_<5 × 10^-4^, AD-stat_2seg, normal_=20.97; μ=0.36, σ=0.30, *p*_2seg, normal_<7.8 × 10^-3^, AD-stat_2seg, normal_=0.99; **2-segment, Weibull**: k=0.50, λ=0.25, *p*_2seg, Weibull_<5 × 10^-4^ , AD-stat_2seg, Weibull_=1.45; k=0.50, λ=0.75, *p*_2seg, Weibull_=0.01, AD-stat_2seg, Weibull_=0.95). These results support the notion that the 2-segment model, which supports an areal gradient of functional properties, better explains the global gradients observed along the dDLPFC (**Figure 7**).

## 3.2 Population-level functional differences along the anterior-posterior axis

To assess whether the five different DLPFC regions exhibit functional differences at the population level, we assigned neurons to regions based on the functional parcellation boundaries derived from the single-neuron gradient analyses performed on all recorded regions, includes areas 46d and 9/46v recorded from Monkey 1 only (**Figure 2G**, **Figure 3 - figure supplement 1, Figure 4 - figure supplement 1**). Boundaries were defined as the median of all significant functional measures that display a functional boundary between both regions (see **Methods**, neuron counts per region are available in **Figure 8 - table supplement 1**). We used the estimated boundaries across all measures with significant 2-segment fits to determine the functional parcellations (including 3-segment models fitted with 46d). This gave the following breaks for Monkey 1: 8Ad-to-9/46d (2.90, shift +0.10), 9/46d-to-46d (9.50, shift +0.00), 8Av-to-9/46v (3.00, shift +0.20). For Monkey 2, 8Av-to-9/46v (1.90, shift −0.90).

To assess the information content at the population level, we used linear discriminant analysis to create cross-temporal decoding plots, where the y-axis shows the time bins used to train the decoders, and the x-axis shows the time bins used to test the decoders (**Figure 8A**). For constructing the decoders we used 100ms time bins with 50ms overlap. Here, we use the term “information” to refer to the classification performance of the decoders. In each plot, we decoded the target or distractor location, separately for correct trials and error trials (**Figures 8-10**). Using these plots, we assessed three aspects of the information contained in each region: 1) information quantity within a period, 2) code stability within a period and 3) code stability across periods. For target and distractor presentation periods, we analyzed the first two measures. For delays 1 and 2, we analyzed all three measures.

**Figure 8.**
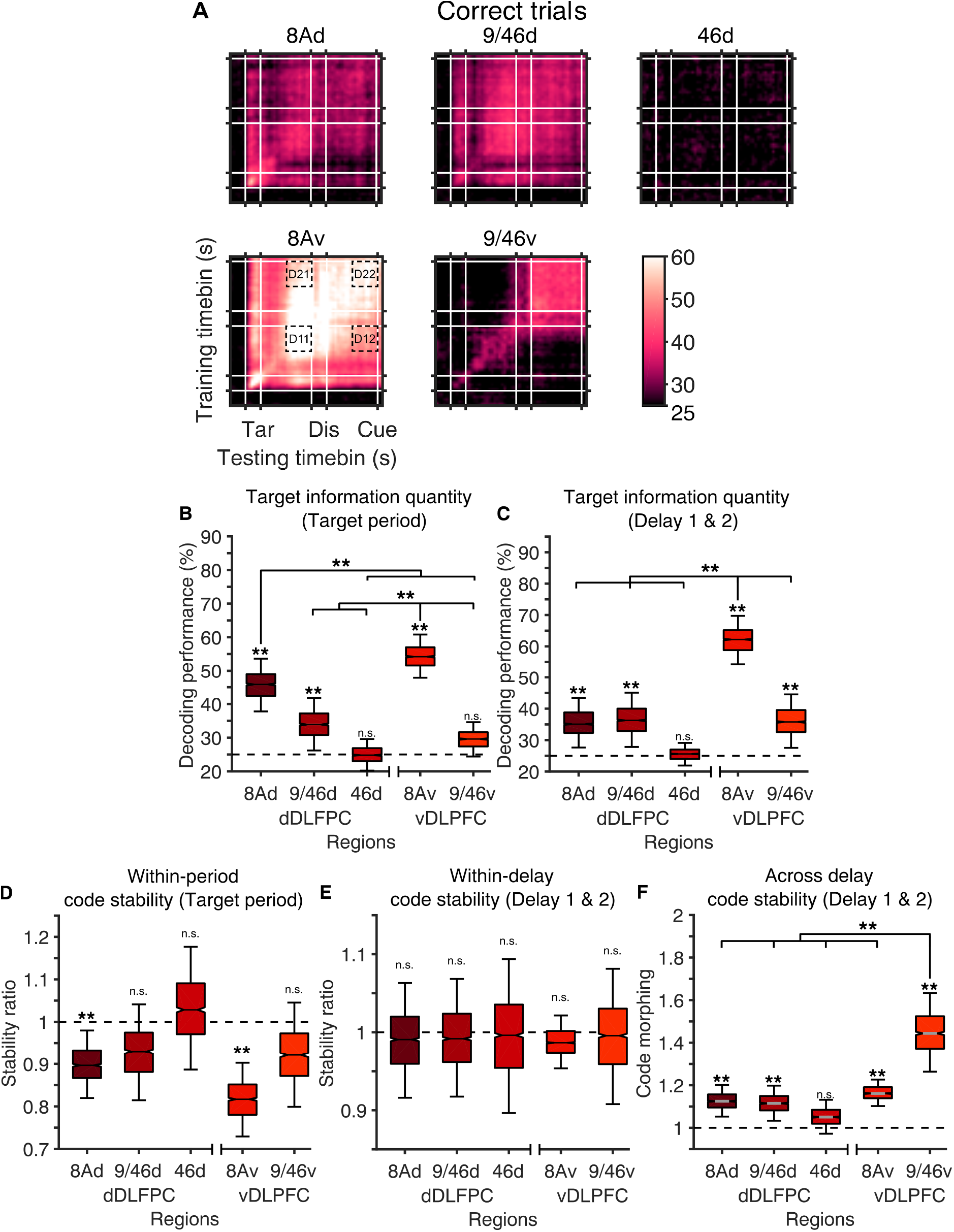
Decoding of the target location. A. , Cross temporal decoding of 4 target locations for 5 DLPFC subregions, **B**, Information quantity during target presentation period, quantified as the performance along the diagonal of the target presentation period, **C**, Information quantity during Delay 1 and Delay 2 periods (average of both delays), quantified as the performance along the diagonal of D11 and D22, **D**, Within-period code stability during the target presentation period, quantified as the ratio of the average performance during target presentation period (without the diagonal), to the diagonal, **E**, Within-period code stability during the delay periods, quantified as the ratio of the performance during the delay period (D11 and D12, without the diagonal), to the diagonal (average of both delays), **F**, Across-delay code stability, quantified as the ratio of decoding performance of D11 to D12 (train in Delay 1, test in Delay 2), and D22 to D21 (average of both ratios).

To compare information quantity and stability across regions we matched the number of locations decoded and the numbers of cells used in each region (n_locations_=4, n_neurons_=28). As area 46d had the minimum number of neurons, we used all 28 neurons in area 46d (lowest number of neurons across regions, **Figure 8 - table supplement 1**), and randomly subsampled 28 neurons for other regions per decoding iteration (n=1000). Since errors were not homogeneously distributed across target locations, we carried out all decoding analyses on the 4 (of 8) target locations that contained a sufficient number of errors (minimum of 6 error trials per session per location) to enable meaningful decoding results in error trials (mean_error trials_=71, range_error trials_=34-113).

### 3.2.1 Target information quantity and stability

From the cross-temporal decoding plots (**Figure 8A**) we first calculated the information quantity, which was the average decoding performance of the 300ms of the target presentation period (**Figure 8B**), or the last 500ms of Delay 1 and Delay 2 (**Figure 8D**, see the same plot but with delay 1 and 2 separated in **Figure 8 - figure supplement 1**), of a time-specific decoder (the diagonal of the plot). We averaged across both delays as they were not significantly different from each other (the 2.5th to 97.5th percentile of the distributions of Delay 1 and Delay 2 values overlapped).

During the target presentation period, we found significant target information in posterior regions in both dorsal and ventral DLPFC (8Ad, 9/46d, 8Av). The information significantly decreased in anterior regions (**Figure 8B**). Decoding was at chance in the most anterior portion of the dDLPFC, area 46d, and the most anterior portion of the vDLPFC, area 9/46v (**Figure 8B**).

During the delay periods, we found significant target information in posterior regions in both dorsal and ventral DLPFC (8Ad, 9/46d, 8Av, 9/46v). In the vDLPFC regions we saw a lower decoding performance in the more anterior region, area 9/46v, however, this trend was not observed in the dDLPFC (**Figure 8C**). Decoding was at chance in the most anterior portion of the dDLPFC, area 46d (**Figure 8C**). We also found that the ventral area 8Av had significantly higher information than any other region analyzed (**Figure 8C**).

We then calculated the within-period code stability, defined as the ratio of the ‘square’ (without the ‘diagonal’) to the ‘diagonal’ of the 300ms of the target presentation period (**Figure 8D**), or the last 500ms of Delay 1 (D11) and Delay 2 (D22) periods (**Figure 8E**). We averaged across both delays as they were not significantly different from each other (95^th^ percentile overlap test, *p*<0.01). A stable code would lead to a ratio close to 1, whereas a dynamic code would lead to a ratio close to 0 (i.e. time-varying code). During the target presentation period, we found a dynamic code in areas 8Ad and 8Av only (**Figure 8D**). However, these dynamic codes were not significantly more dynamic than the codes found in more anterior regions (**Figure 8D**). During the delay periods, we found that all regions have stable within-delay codes, as their ratios were not significantly different from 1 (**Figure 8E**).

Finally, we calculated the across-delay code stability. A lack of across-delay code stability is referred to as code-morphing (Parthasarathy et al., 2017). Code-morphing is present when the neuronal code in Delay 1 can be used to predict the target location in Delay 1 but cannot be used to predict the target location in Delay 2, and vice versa. To calculate the across-delay code stability, we calculated the ratio between the performance of a decoder trained and tested in the last 500ms of Delay 1 (D11 ‘square’, **Figure 8A**), and a decoder trained in Delay 1 but tested in Delay 2 (D12 ‘square’, **Figure 8A**), i.e. D11/D12. We did the same for Delay 2, i.e. D22/D21. A code that is perfectly stable across delays (i.e. that does not morph) would lead to a Delay 1 and Delay 2 code-morphing value of 1 (i.e. D11=D12 and D22=D21), while a code that morphs would have a value larger than 1 for both Delay 1 and Delay 2 (i.e. D11>D12 and D22>D21). We found code-morphing in all regions except for area 46d (likely due to the low information content present in 46d in the first place). Along the anterior-posterior axis, we observed a significant increase in code-morphing from the posterior 8Av to the anterior region 9/46v (**Figure 8F**).

### 3.2.3 Distractor information quantity and stability

We previously showed that posterior regions of the dDLPFC have a higher proportion of cells selective to the distractor location and also a higher proportion of cells that filter the distractor (**Figure 3D, H**). To determine how these single-neuron properties relate to information about distractor location at the population level, we trained the decoder to predict the location of the distractor, rather than the target (**Figure 9**).

**Figure 9.**
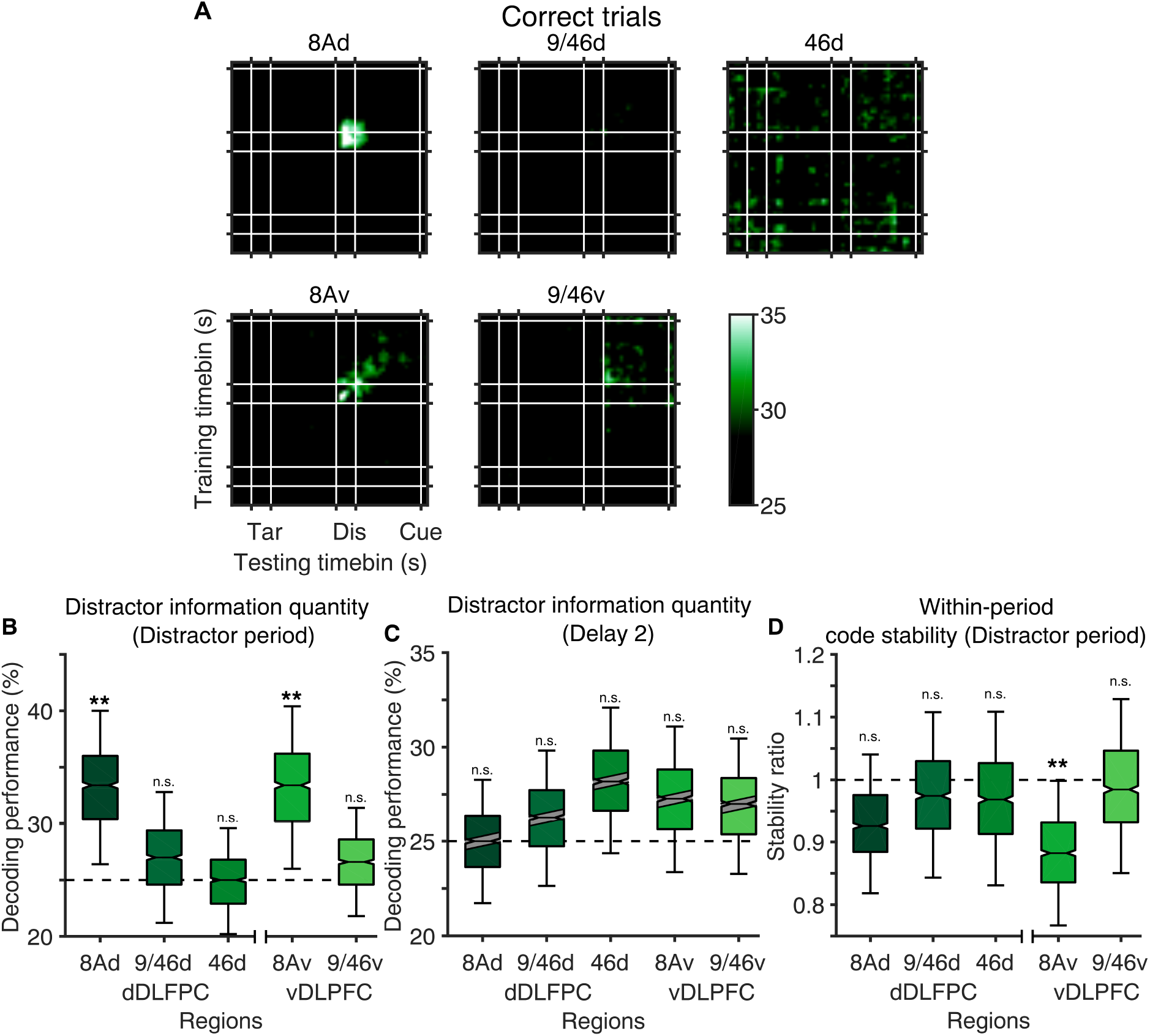
Decoding of the distractor location. **A**, Cross temporal decoding for 5 subregions, **B**, Information quantity in Delay 2 for the above regions **C**, Information stability in Delay 2 for the above regions. The period for comparing correct and error trials was defined as the last 500ms of Delay 2. **D**, Within-period code stability during the distractor presentation period, quantified as the ratio of the average performance during the distractor presentation period (without the diagonal), to the diagonal.

During the distractor presentation period, we found significant distractor information in posterior regions in both dorsal and ventral DLPFC (8Ad, 8Av). The information in posterior regions was not significantly different from that found in anterior regions (**Figure 9B**). Decoding was at chance in the most anterior portions of the dDLPFC, areas 9/46d, and 46d, and the most anterior portion of the vDLPFC, area 9/46v (**Figure 9B**).

During the Delay 2 period, we found that none of the regions contained significant distractor information (**Figure 9D**).

During the distractor presentation period, we found a dynamic code in area 8Av (**Figure 9C**). However, this dynamic code was not significantly more dynamic than the codes found in other regions (**Figure 9C**).

### 3.2.4 Information quantity and stability in error trials

To assess the behavioral relevance of the activity of different regions, we calculated the decoding performance in error trials and compared it with the decoding performance in correct trials (**Figure 10A**). Error trials are those in which the monkey waited for the go-cue to respond, but responded to an incorrect location, or failed to initiate a response within 1 second. Behavioral relevance would be reflected as a significant difference between the decoding performance of correct and error trials.

**Figure 10.**
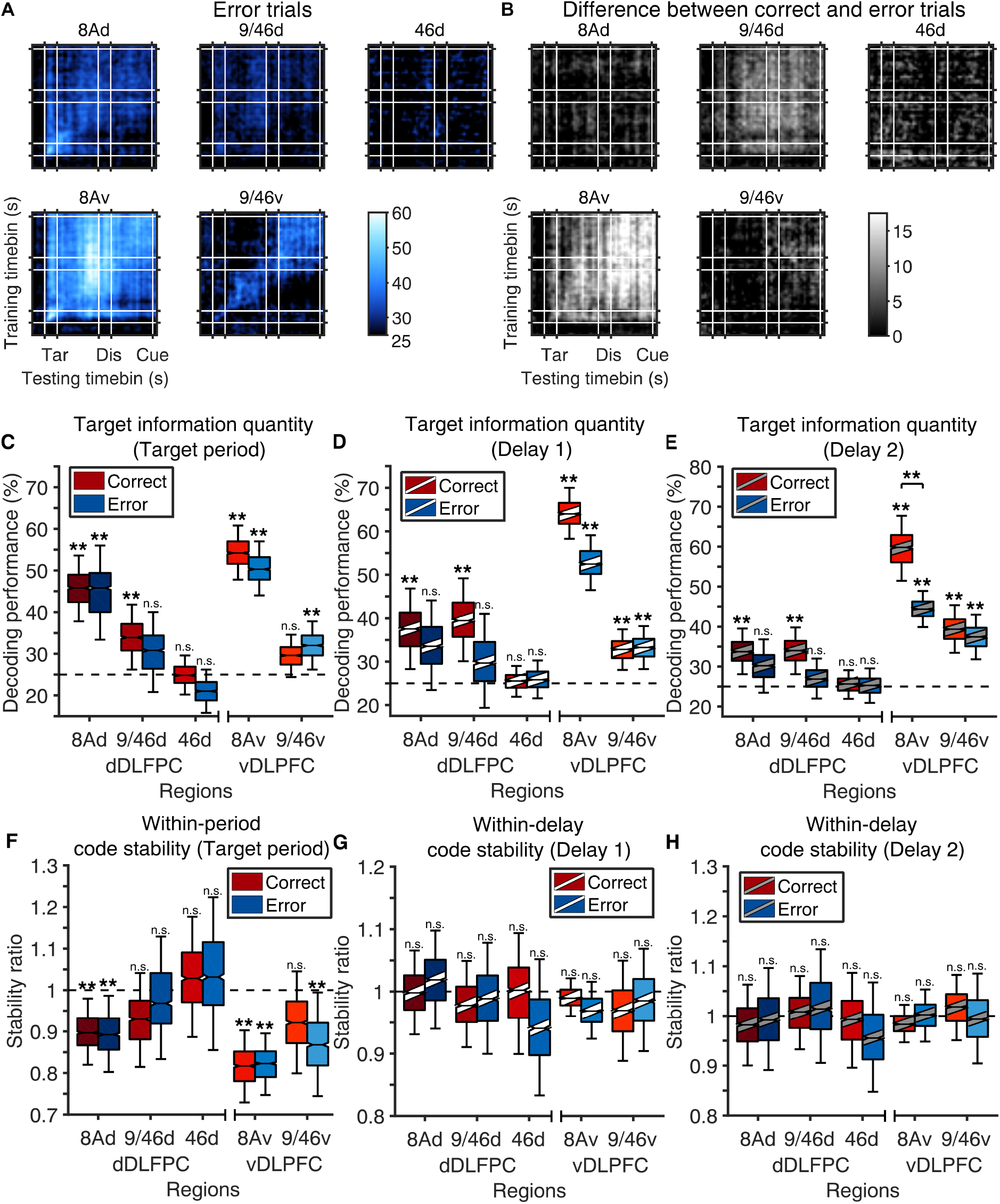
Decoding of the target location in correct vs error trials. **A**, Cross temporal decoding for 5 subregions in error trials, **B**, Same as A, but shows the difference between correct and error trials, **C-E**, Information quantity in the target period, delay 1 (white stripe) and delay 2 (grey stripe) for the above regions in correct (red) and error (blue) trials, **F-G**, Information stability in target period, delay 1 (white stripe) and delay 2 (grey stripe) for the above regions.

The only region to show a difference in information between correct and error trials was area 8Av, which had less target information in error trials during the Delay 2 period (**Figure 10E**). None of the regions showed significant differences in the target period or the Delay 1 period information, nor in code stability (**Figure 10**, **Figure 10 - figure supplement 1**). Furthermore, none of the regions showed a difference between correct and error trials in the encoding of distractor information, suggesting that errors were not driven by increased distraction (**Figure 10 - figure supplement 2**).

## Discussion

Here we show that the functional gradient in the DLPFC is the result of functionally-distinct areas organized as a gradient, rather than a smooth gradient of functional properties along the anterior-posterior axis. We showed this by analyzing the response properties of single neurons along the anterior-posterior axis (**Figures 3-6**), and further dissected the differences between regions at the population level (**Figures 8-10**).

### 4.1 Global functional gradients are the result of functionally-distinct areas organized as a gradient

Previous studies have described the existence of global functional gradients in the DLPFC (Riley et al., 2018, 2017). In line with this, we found global functional gradients, including posterior to anterior decreases in selectivity and memory field sizes during stimulus and delay periods, distractor filtering, and mixed selectivity, and increases in latencies. While it is challenging to match our microelectrode positions to those reported in the relevant studies by Riley and colleagues, we estimate that our recording electrodes in both the dDLPFC and vDLPFC fall within what they refer to as the “dorsal DLPFC” (i.e. regions dorsal to the principal sulcus, as well as the ventral lip of the principal sulcus). With that consideration, the global gradients we report here are largely consistent with those reported previously (Riley et al., 2018, 2017).

However, we extend these studies by showing that these global gradients are the result of functionally-distinct areas organized as a gradient, rather than a smooth gradient of functional properties. To our knowledge, ours is the first study to explicitly assess whether the functional heterogeneity observed at the cellular level in the DLPFC can be better explained by a smooth gradient of functional properties, or by functionally-distinct areas organized as a gradient. Our results support the latter interpretation.

### 4.2 A prominent role of posterior DLPFC in the maintenance of spatial working memory information

The following lines of evidence suggest that the posterior DLPFC regions play a special role in the maintenance of spatial working memory information. We found that area 8Av had more target information than other regions (**Figure 8**). Area 8Av was the only region that showed differences in decoding performance between correct and error trials, having a lower decoding performance in error trials during Delay 2 (**Figure 10**). Area 8Av also has the greatest proportion of nonlinear mixed selective cells of the regions recorded (**Figure 8 - figure supplement 3**). This observation is consistent with a previous study that showed that optogenetic inactivation of the frontal eye fields (which is the posterior aspect of what we categorized here as area 8A) during the delay period of a memory-guided saccade task leads to a decrease in task performance (Acker et al., 2016). Overall, these results suggest that among DLPFC regions, posterior regions, and in particular area 8Av, play a special role in the maintenance of spatial working memory information.

### 4.2 A prominent lack of involvement of anterior dDLPFC in the maintenance of spatial working memory information

An unexpected observation was the almost absolute absence of target information in the most anterior region analyzed (area 46d). In this region, very few cells were selective (**Figure 3, Figure 3 - figure supplement 1, Figure 4, Figure 4 - figure supplement 1**), and we could not decode target information above chance (**Figure 8, Figure 8 - figure supplement 1 and 2**). This observation suggests that spatial memory processing is restricted to the posterior DLPFC, without the involvement of anterior regions.

### 4.3 Code stability

The within-delay information stability, which is the ratio of the off-diagonal ‘square’ to the ‘diagonal’ in the last 500ms of each delay, was close to 1 across all regions (**Figure 9C**). This suggests that there is strong stability of the code within delay periods across the DLPFC. On the other hand, the across-delay stability, which is a measure of how generalizable is the memory code across delays, was lower in the vDLFPC, and in particular in the anterior region 9/46v (**Figure 9D**). This observation may be explained by the higher proportion of nonlinear mixed selectivity cells in vDLPFC compared to dDLPFC (Monkey 1: mean_dDLPFC_ = 7.4%, mean_vDLPFC_ = 36.5%, z_vDLPFC>dDLPFC_ = 5.79, *p*_vDLPFC>dDLPFC_ = 6.86 × 10^-9^, 2-tailed Mann-Whitney U test; Monkey 2: mean_dDLPFC_ = 21.6%, mean_vDLPFC_ = 53.6%, z_vDLPFC>dDLPFC_ = 3.76, *p*_vDLPFC>dDLPFC_=1.73 × 10^-4^, 2-tailed Mann-Whitney U test, **Figure 8 - figure supplement 3**), since non-linear mixed selectivity plays a large role in code-morphing in the DLPFC (Parthasarathy et al., 2017).

### 4.4 Conclusion

Overall, our results support the notion that functionally-dissociable areas in the DLPFC are organized along the anterior-posterior axis in a functional gradient. This type of organization has been observed in other brain systems, such as the visual system, where individual regions (V1, V2, V4, IT), which are anatomically and functionally distinct from each other, are arranged along an anterior-posterior functional gradient, with more anterior regions having larger receptive fields and more complex response selectivities (Freud et al., 2017; Hubel and Wiesel, 1959; Lerner, 2001; O’Rawe and Leung, 2020; Tsao et al., 2006). The ability to record large numbers of neurons simultaneously across multiple brain regions in behaving animals provides an important resource for systems neuroscience research (Dotson et al., 2018; Mitz et al., 2017). Crucially, our results underscore the importance of respecting the functional boundaries between regions when analyzing these large datasets, since it is common practice to group cells from multiple regions for analyses (Bartolo et al., 2020; Parthasarathy et al., 2019, 2017; Tang et al., 2020). These results will also be important for researchers using artificial neural networks to model cognitive processes, since a single “prefrontal module”, usually modelled as a randomly-connected recurrent neural network or as a bump-attractor network, may not reflect functional distinctions and complex interactions between different prefrontal areas (Mante et al., 2013; Parthasarathy et al., 2019; Tang et al., 2020; Wimmer et al., 2014; Yang et al., 2019).

## Methods

### 5.1 Animals and single-unit recordings

We used two male adult macaques *(Macaca fascicularis)* in the experiments: Monkey 1 (age 4) and Monkey 2 (age 6). All animal procedures were approved by and conducted in compliance with the standards of the Agri-Food and Veterinary Authority of Singapore and the Singapore Health Services Institutional Animal Care and Use Committee (SingHealth IACUC #2012/SHS/757). Procedures also conformed to the recommendations described in Guidelines for the Care and Use of Mammals in Neuroscience and Behavioral Research (Van Sluyters and Obernier, 2003). We first implanted a titanium head-post (Crist Instruments, MD, USA), followed by intracortical microelectrode arrays (MicroProbes, MD, USA) in the left frontal cortex. In Monkey 1, we implanted 6 arrays of 16 electrodes, 1 array of 32 electrodes, and 2 arrays of 32 electrodes for a total of 192 electrodes. In Monkey 2, we implanted 1 array of 16 electrodes, 2 arrays of 32 electrodes, and 2 arrays of 16 electrodes in the cortex for a total of 112 electrodes. For additional details on the surgery refer to Parthasarathy et al., 2017 (Parthasarathy et al., 2017).

### 5.2 Recording techniques

Neural signals were acquired using a 128-channel and a 256-channel Plexon OmniPlex system (Plexon Inc, TX) at a sampling rate of 40 kHz. Wide-band signals were band-pass filtered between 250 and 10,000 Hz. Subsequently, spikes were detected using an automated Hidden Markov-Model-based algorithm for each channel (Herbst et al., 2008). Eye positions were obtained with an infrared-based eye-tracking device from SR Research Ltd. (Eyelink 1000 Plus). The behavioral task was designed on a standalone PC (stimulus PC) using the Psychophysics Toolbox in MATLAB (Mathworks, MA)(Brainard, 1997). To align the neural and behavioral activity (trial epochs and eye data) for downstream data analysis, we employed strobe words denoting trial epochs and performance (rewarded or failure) during the trial. These strobe words were generated on the stimulus PC and sent to the Plexon and Eyelink computers via the parallel port.

### 5.3 Task

Each trial started with a mandatory period (500ms) during which the animal fixated on a white circle at the center of the screen. While continuing to fixate, the animal was presented with a target (a red square) for 300ms at any one of eight locations in a 3 × 3 grid. The center square of the 3 × 3 grid contained the fixation spot and was not used. The presentation of the target was followed by a delay of 1,000ms, during which the animal was expected to maintain fixation on the white circle at the center. At the end of this delay, a distractor (a green square) was presented for 300ms at any one of the seven locations (other than where the target was presented). This was again followed by a delay of 1,000ms. At the end of the second delay, the animal was given a go-cue (the disappearance of the fixation spot) to make a saccade towards the target location presented earlier in the trial. Saccades to the target location within 150 ms and continued fixation at the saccade location for 200ms was considered a correct trial. An illustration of the task is shown in **Figure 1A**. Because of a behavioral bias in one of the animals, the target was presented in only seven of the eight target locations in that animal (we excluded the location at the bottom-right).

### 5.4 Estimation of electrode positions

We calculated the anterior-posterior location of each electrode using photos taken during the implantation surgeries. We used the electrode arrays, which have a fixed size, to convert pixels to millimeters. The posterior tip of the principal sulcus was used as the reference, such that positive values denote more anterior positions, and negative values denote more posterior positions (**Figure 1A**).

### 5.5 Estimation of anatomical parcellations for each monkey

We estimated the anatomical boundaries of each region from a magnetic resonance imaging atlas of the cynomolgus monkey brain in native millimeter space (Frey et al., 2011). The anatomical boundaries identified from the atlas had the following values: 8Ad-9/46d (2.8mm), 9/46d-46d (9.5mm), 8Av-9/46v (3.0mm), 9/46V-46v (9.7mm).

### 5.6 Calculating functional measures per electrode

To determine if a neuron was selective during the target, distractor, and/or delay periods, we averaged the spike count across the entire period (target period: 0-0.3s; delay 1: 0.3-1.3s; distractor: 1.3-1.6s; delay 2: 1.6-2.6s) and run a 1-way ANOVA across locations (threshold of *p*<0.05).

To calculate response latencies, we identified the earliest responsive bin for the preferred location for each neuron across the entire trial in 100ms bins with 50ms overlap (significantly above the pre-stimulus baseline of −300ms to 0ms; t-test, *p*<0.05). If there were no preferred locations, we identified the earliest responsive bin across all locations.

To calculate receptive field size and memory field sizes, we counted the number of target locations where activity during the period analyzed (target, Delay 1 or Delay 2) was significantly different from the pre-stimulus baseline (−300ms to 0ms).

To calculate selectivity indices during the target presentation or delay periods, we calculated the index as *(max-min)/(max+min)*, where max is the average firing rate for the target location with the highest firing rate and min is the corresponding location with the lowest firing rate (min).

To calculate distractor filtering, we obtained the ratio between the average firing rate for the preferred location during the target presentation period, to the firing rate for the same distractor location during the distractor presentation period.

To determine if a neuron had non-linear mixed selectivity we measured the average activity of the last 500ms of Delay 1 and Delay 2, and carried out a 2-way ANOVA, using target location, and epoch (Delay 1 or Delay 2) as the two factors. Neurons that showed a significant interaction term (with a threshold of *p*<0.05) were considered non-linear mixed selective.

### 5.7 Fitting the linear (1-segment) and piecewise (2- and 3-segment) models

For each of the 13 functional measures assessed, single neuron data was first smoothed along the AP-axis using a 0.5mm window, with 0.17mm overlap (i.e. 1/3 of window size). To describe the trend along the anterior-posterior axis, we first fitted a 1-segment, 2-segment, and 3-segment (only applicable in Monkey 1) robust linear regression model to the smoothed dataset to reduce the bias from outlier datapoints (MATLAB function fitlm(), with the ‘Robust’ parameter set to ‘on’. This robust linear fitting uses the default ‘bisquare’ weighting function). To quantify the goodness-of-fit of a given functional measure, we calculated an adjusted R^2^ for the linear fit of the actual dataset, which penalizes the number of parameters used in the model, and thus penalizes the increasing number of segments fitted per model:

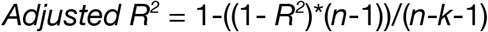

where *R^2^* is the R^2^ calculated using the robust regression, *n* is the number of data points used for model fitting, and *k* is the number of independent regressors of the fitted model (the number of variables in the model)(Cohen et al., 2003).

For the 2-segment models, we first defined the possible break locations, which were ±1.5mm from the estimated anatomical boundaries (between areas 8Ad-9/46d, and 8Av-9/46v). This ±1.5mm area was subdivided into 30 non-overlapping 0.1mm steps, and we fitted a line to the left and right of these breaks using robust linear regression. We imposed that all lines require at least 3 data points to be included in the analysis. Out of the 30 models, the model with the lowest root mean squared error (RMSE) was considered to be optimal, and thus its break location was used as the functional boundary for that functional measure. For the 3-segment models with two breakpoints (only for Monkey 1’s dDLPFC), the procedure was identical, except that both breaks were fit sequentially from posterior to anterior. For instance, we first identified the break between 8Ad and 9/46d), and we then identified the second break between 9/46d and 46d. If two models were tied with similar breakpoints (e.g. both at the boundary between 8Ad and 9/46d), we picked the model with the breakpoint closest to the estimated anatomical boundary. The 2- and 3-segment plots in the figures use the best breakpoints identified in this manner.

Note that some of the functional measures could not be calculated for all cells (i.e. distractor filtering, which only used cells with significant target selectivity). As such, for all 13 functional measures, we only used the subset of cells for which all measures could be calculated (569 out of 632). This preserves the number of data points used for gradient fitting and avoids a bias from varying numbers of datapoints per measure.

### 5.8 Testing for the presence of global gradients

To assess whether there was a global gradient along the anterior-posterior axis, we tested whether the adjusted R^2^ of the 1-segment fit was significant compared to a shuffled distribution of adjusted R^2^. Specifically, we obtained a shuffled distribution by shuffling the dataset (post-smoothing) along the AP-axis and fitting the same robust linear model to these shuffled datasets for 1000-iterations. If the adjusted R^2^ of the actual fit was larger than the 95^th^ percentile of the shuffled distribution we considered it significant and determined that a global gradient was present (displayed in figures as an adjusted R^2^ in bold font above the 1-segment models). This was done separately for the dDLPFC and vDLPFC, and independently for each monkey. We corrected for multiple comparisons with the FDR Benjamini-Hochberg method, separately for the dDLPFC and vDLPFC of each monkey as they were analyzed for gradients independently.

### 5.8 Testing for whether the data was better described by a smooth gradient or an areal gradient

To test whether the 1-segment model (which would support the smooth gradient hypothesis) fitted the data better than the 2-seg model (which would support the areal gradient hypothesis), we compared the 1-segment model’s adjusted R^2^ to a null distribution of 1-segment adjusted R^2^s generated from surrogate datasets (1000 iterations). These surrogate datasets had mean and variance within 30% of the actual data, and, importantly, had a *2-segment* adjusted R^2^ within a fixed 20% of the actual 2-seg model fit (see full statistical flowchart in **Figure 2 – figure supplement 2**). Note that as it was challenging to generate 2-segment surrogate datasets for the nonlinear mixed selectivity % in the vDLPFC of Monkey 1 according to the criterion above, so we relaxed the criterion for this specific measure in the vDLPFC for Monkey 1. Namely, this 2-segment surrogate dataset had mean and variance within 100% of the actual data, and a *2-segment* adjusted R^2^ within a fixed 60% of the actual 2-seg model fit. The distribution of 1-segment adjusted R^2^s obtained from this surrogate dataset represents a null distribution of what would be expected by chance given a dataset with that specific 2-segment model fit. If the actual 1-segment model fit exceeded the 95^th^ percentile of the null distribution, we concluded that the 1-segment model fitted the data better than the 2-segment model. To test whether the 2-segment model fitted the data better than the 1-seg model, we followed a similar procedure, except that the surrogate data matched the 1-segment adjusted R^2^, and the null distribution was a 2-segment adjusted R2 distribution. A similar procedure was used to compare the 1-segment and 3-segment models, and 3-segment and 1-segment models.

Next, for all the functional measures with a significant global gradient, we tested if the distributions of 1-segment and 2-segment model-comparison *p*-values followed a Weibull distribution biased towards zero using the Anderson-Darling test (see **Figure 7**). To this end, we tested 2 alternative hypotheses: (1) that the 1 - or 2-segment distributions follow a Weibull distribution biased towards zero, or (2) that the 1- or 2-segment distributions follow a normal distribution with μ being equal to the mean of each distribution. The logic of this analysis is that if the global gradients are better explained by an areal gradient (i.e. are better fitted by a 2-segment distribution) we expect to see the model-comparison *p*-values of the 2-segment distribution following a Weibull distribution biased towards zero, but not the normal distribution described above. Similarly, if the global gradients are better explained by a smooth gradient (i.e. are better fitted by a 1-segment distribution) we expect to see the model-comparison *p*-values of the 1-segment distribution following a Weibull distribution biased towards zero, but not a normal distribution.

### 5.9 Estimation of functional parcellations for each monkey

To identify the functional parcellation boundaries for each animal, we collated the boundaries obtained from all the models that showed a significant 2- or 3-segment fit. The median of all these points was the final estimated functional boundary for those regions. Each monkey’s final estimated parcellation map was used to split neurons into regions depending on their electrode positions. These neurons were then used in the population decoding analyses.

### 5.11 Cross-temporal decoding analysis

We pooled all neurons per region (using the functional boundaries) and subsampled to the minimum number of neurons across regions (n=28, minimum in area 46d). We used data at each time point (100ms bins with 50ms overlap) to train a decoder using Linear Discriminant Analysis (LDA, MATLAB function classify()). We tested the decoder on data from all 100ms time bins in the trial, as described in Parthasarathy et al. 2017 (Parthasarathy et al., 2017). When decoding data in the full space, we denoised the training and testing data using principal components analysis (PCA) at every time point by reconstructing the data with the top *n* principal components that explained at least 90% of the variance. The target locations were predicted by training and testing the decoder on datasets from equivalent time points. The performance of the decoder, a measure of the information about the target location in the population activity, was computed at each timepoint as a percentage of test trials in which the target location was predicted correctly. This process was repeated 1,000 times, with different subsets of correct trials used to constitute the training and test sets. We used 1500 trials for the training set and 100 trials for the test set, for both the correct and error trial analyses. For the supplementary decoding analysis which featured only Monkey 1’s neurons (**Figure 8 – figure supplement 2**), the only deviation from the above protocol is that we used all neurons in each region for that monkey.

### 5.12 Data and code availability statement

The code package and data needed to perform the analyses used in the paper are available at https://github.com/PK-HQ/gradient-dlpfc.

## Acknowledgements

This work was supported by start-up grants from the Ministry of Education Tier 1 Academic Research Fund and SINAPSE to CL, a grant from the Ministry of Education Tier 2 Academic Research Fund to CL (MOE2016-T2-2-117), and a grant from the Ministry of Education Tier 3 Academic Research Fund to CL (MOE2017-T3-1-002). We would like to thank Shih-Cheng Yen for useful comments on the manuscript.Additional information

## Funding

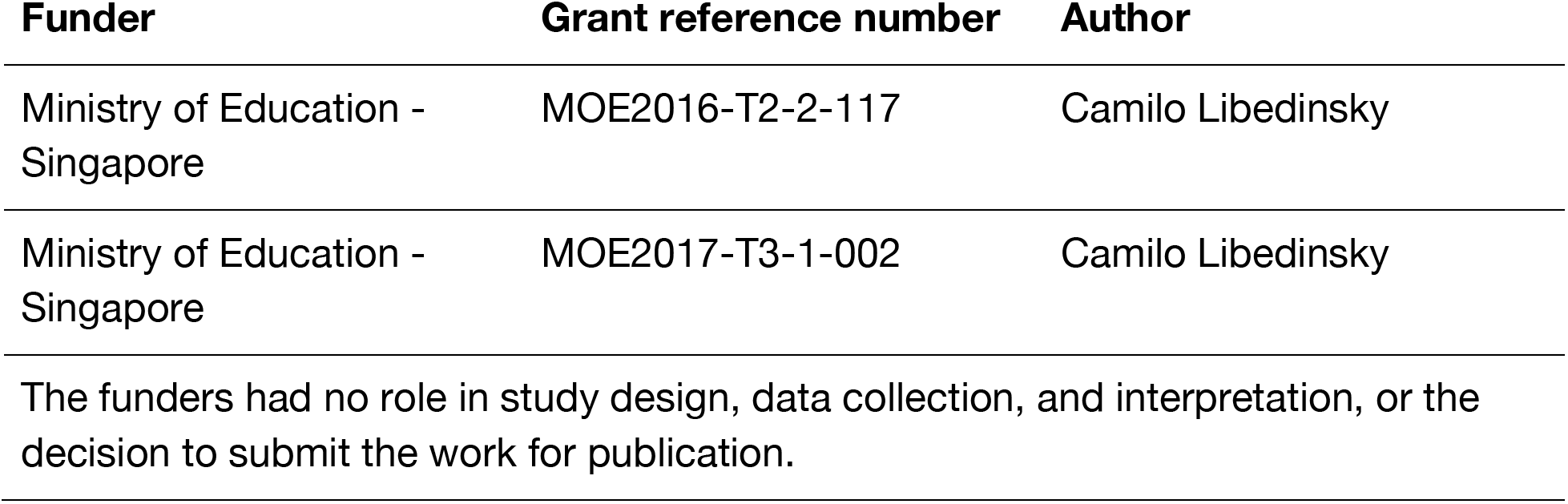

## Author contributions

Pin Kwang, Conceptualization, Data curation, Software, Formal analysis, Investigation, Visualization, Methodology, Writing - original draft, Writing - review and editing; Cheng Tang, Data curation, Software, Formal analysis, Methodology; Roger Herikstad, Data curation, Software, Investigation, Methodology; Arunika Pillay, Conceptualization, Visualization Camilo Libedinsky, Conceptualization, Resources, Supervision, Validation, Investigation, Methodology, Writing - original draft, Project administration, Writing - review and editing

## Ethics

All animal procedures were approved by and conducted in compliance with the standards of the Agri-Food and Veterinary Authority of Singapore and the Singapore Health Services Institutional Animal Care and Use Committee (SingHealth IACUC #2012/SHS/757). The procedures also conformed to the recommendations described in Guidelines for the Care and Use of Mammals in Neuroscience and Behavioral Research(Van Sluyters and Obernier, 2003).

## Figure supplements

**Figure 2 – figure supplement 1.**
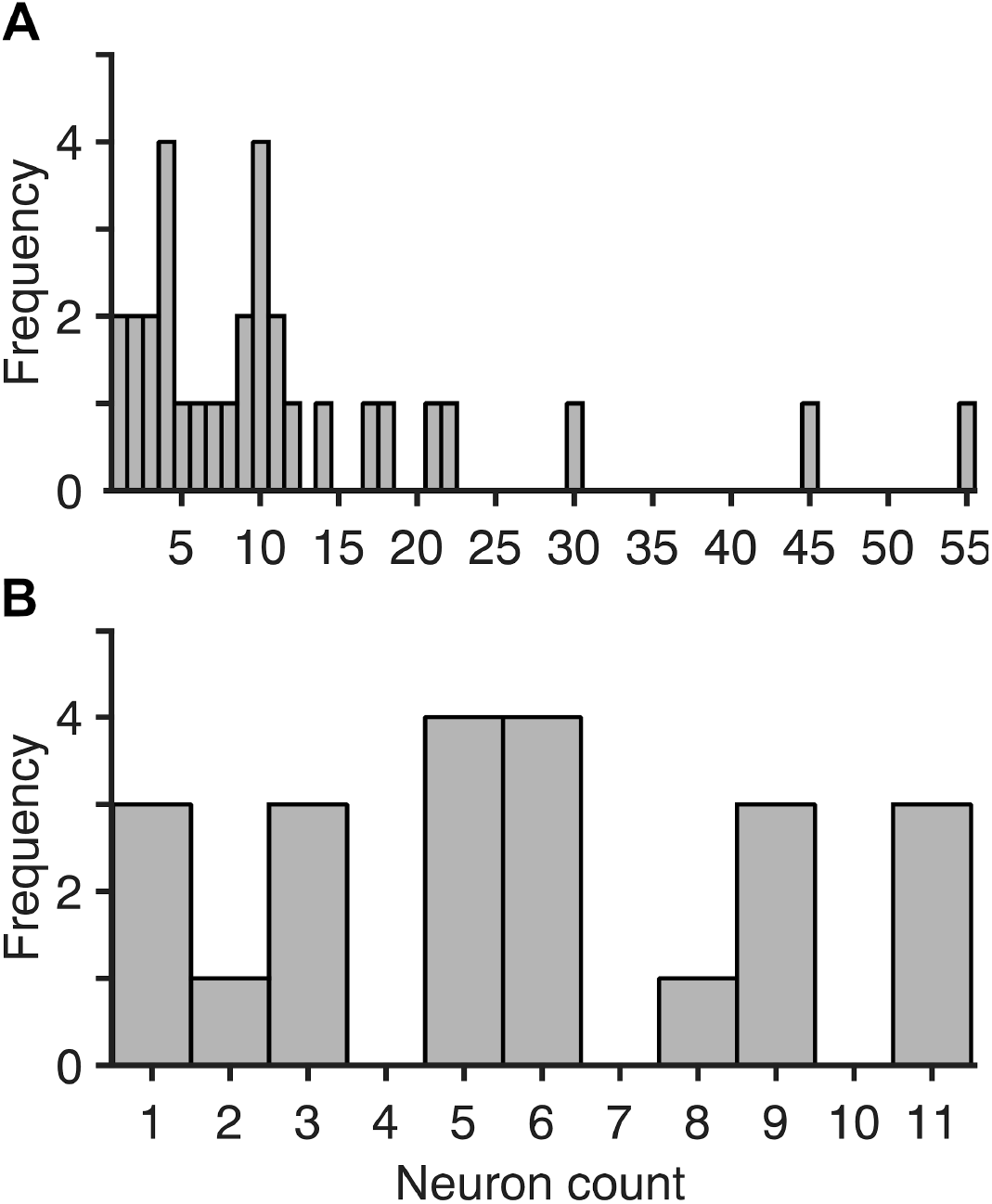
Histogram of frequency by neuron count per data point. **A**, Monkey 1, **B**, Monkey 2.

**Figure 2 – figure supplement 2.**
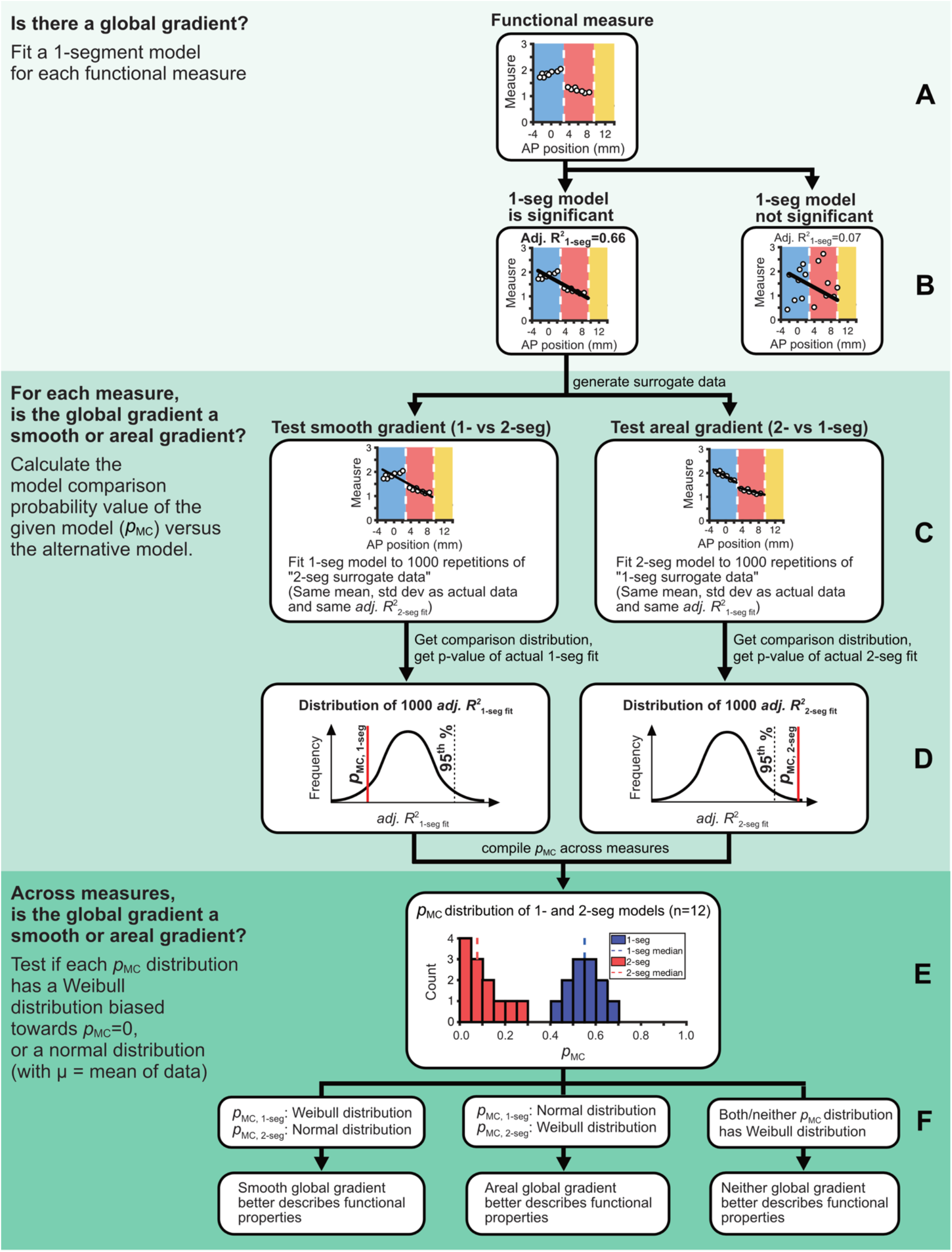
Statistical flowchart for determining the best fitting model for data. Diagram showing the flow of statistical tests used to test whether there is a global gradient and, if there is a global gradient, whether a smooth or areal gradient explains this global gradient better (bottom). **A**, Functional measure along the anterior-posterior axis, **B**, Testing the whether there was a significant global gradient, **C**, Generating 1-segment and 2-segment surrogate datasets, **D**, Obtaining the comparison-distribution of adjusted R^2^, and the comparison-distribution *p*-value (*p*_MC_) for the model tested, **E**, To test if data is skewed towards *p*_MC_=0, we fit normal and Weibull models to the data, **F**, Interpret the normal and Weibull fits of the 1-segment and 2-segment data.

**Figure 3 – figure supplement 1.**
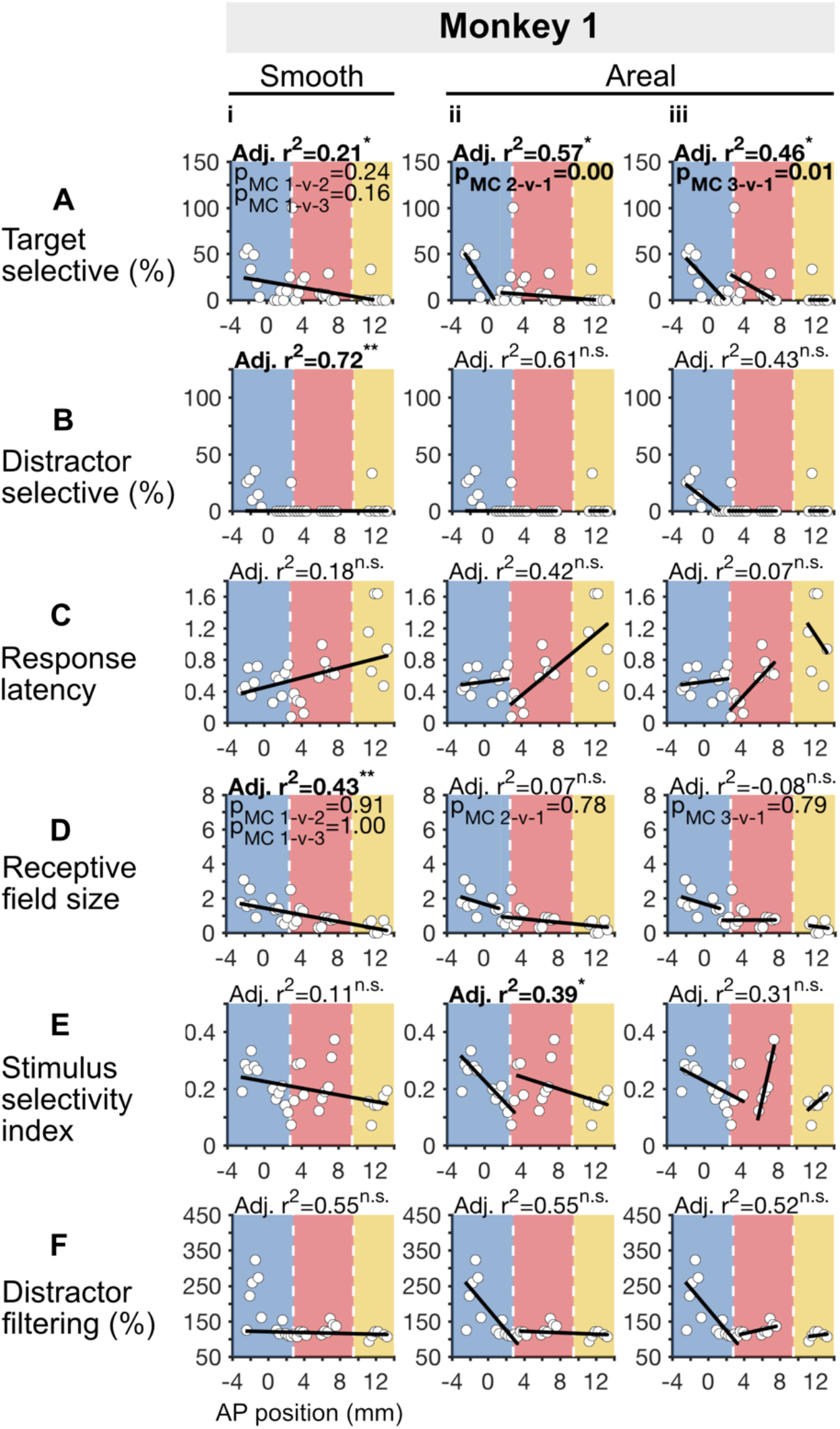
Target period gradient fits but with all 3 segments fit for Monkey 1. Same as **Figure 3**, but with the area 46d datapoints included in model fitting.

**Figure 4 – figure supplement 1.**
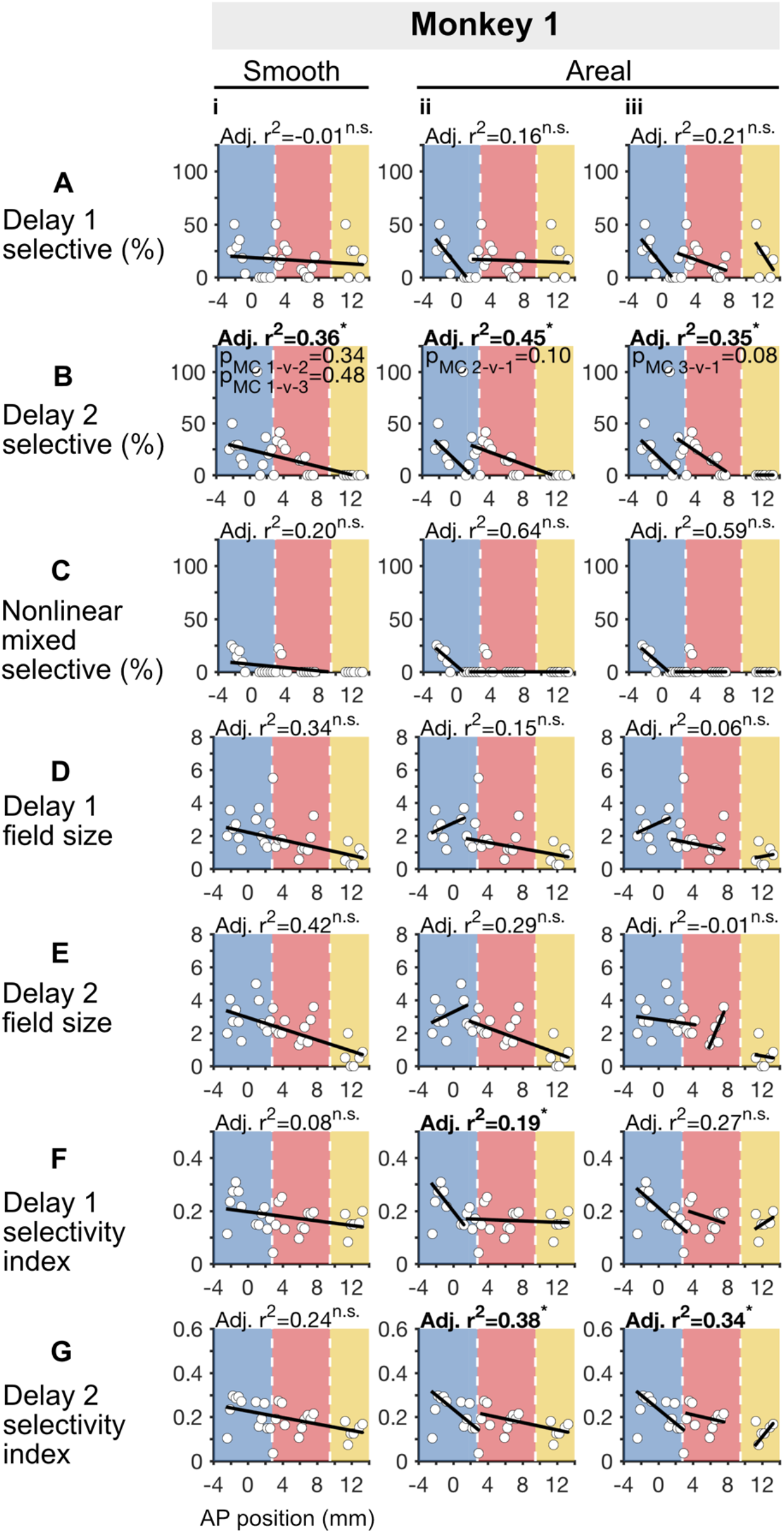
Delay period gradient fits but with all 3 segments fit for Monkey 1. Same as **Figure 4**, but with the area 46d datapoints included in model fitting.

**Figure 7 – figure supplement 1.**
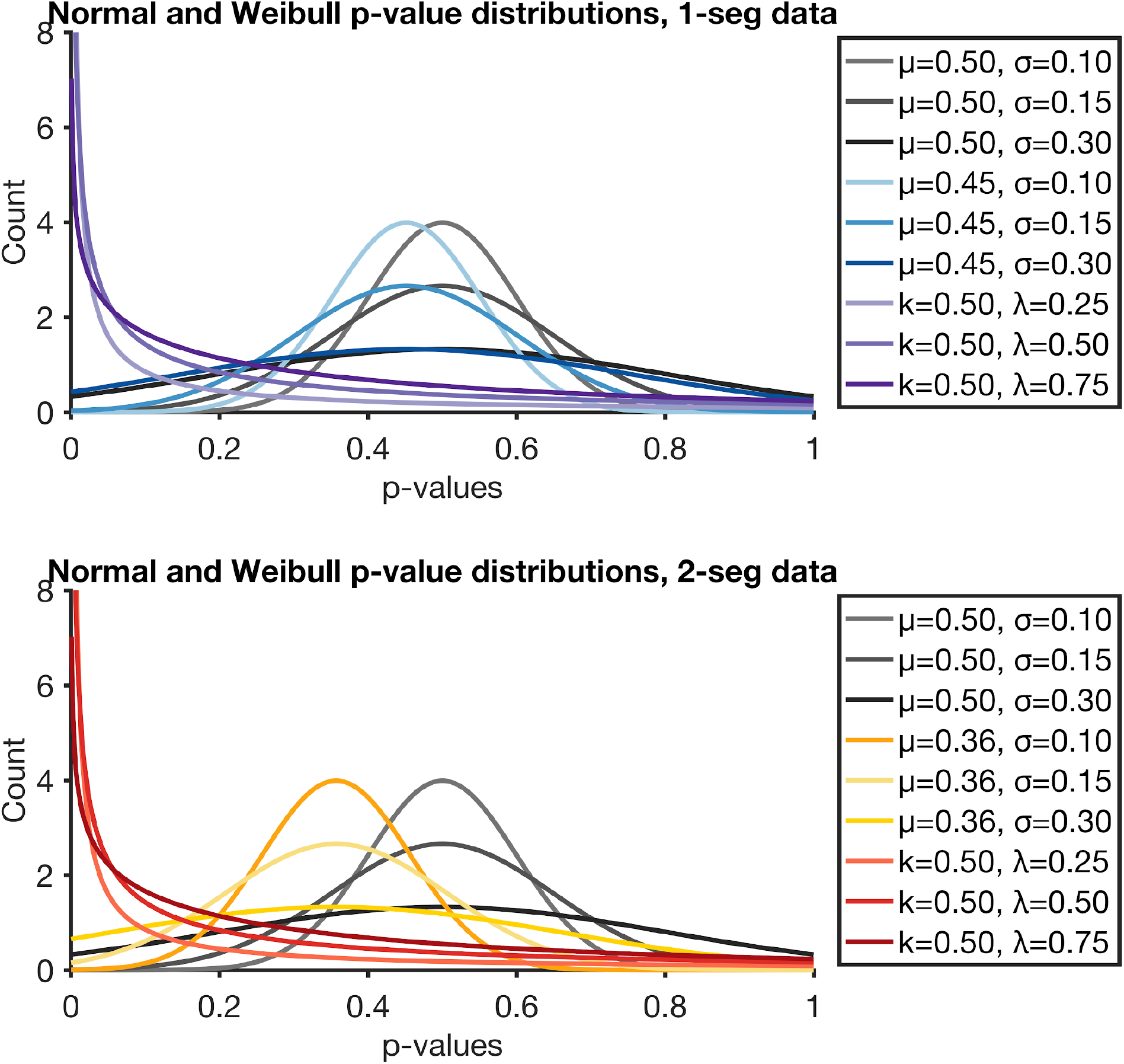
normal and Weibull distributions used for the Anderson Darling test of 1-segment and 2-segment data. Model parameters for each line are shown in the figure legend. k=Weibull shape parameter, λ=Weibull scale parameter, μ=normal mean parameter, σ=normal standard deviation parameter.

**Figure 8 - table supplement 1.** Recorded neurons per region. The number of neurons recorded in each region for each monkey.

| Region | Subregion | Monkey 1 | Monkey 2 |
| --- | --- | --- | --- |
| **Dorsal DLPFC** | 46d | 28 | 0 |
|  | 9/46d | 72 | 52 |
|  | 8Ad | 146 | 22 |
| **Ventral DLPFC** | 9/46v | 50 | 0 |
|  | 8Av | 7 | 112 |

**Figure 8 – figure supplement 1.**
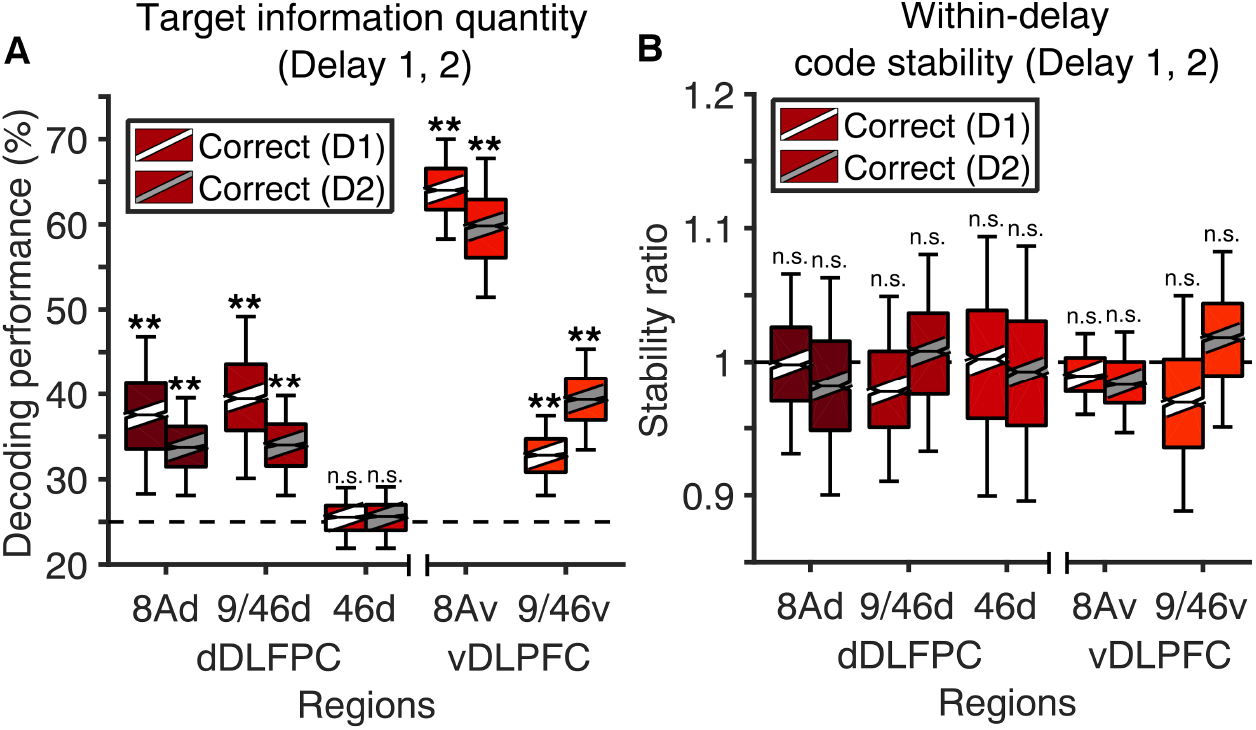
Regional differences in Delay 1 and 2 target information quantity and stability in correct trials. **A**, Information quantity in delays 1 and 2 for the above regions, **B**, Information stability in delays 1 and 2 for the above regions

**Figure 8 – figure supplement 2.**
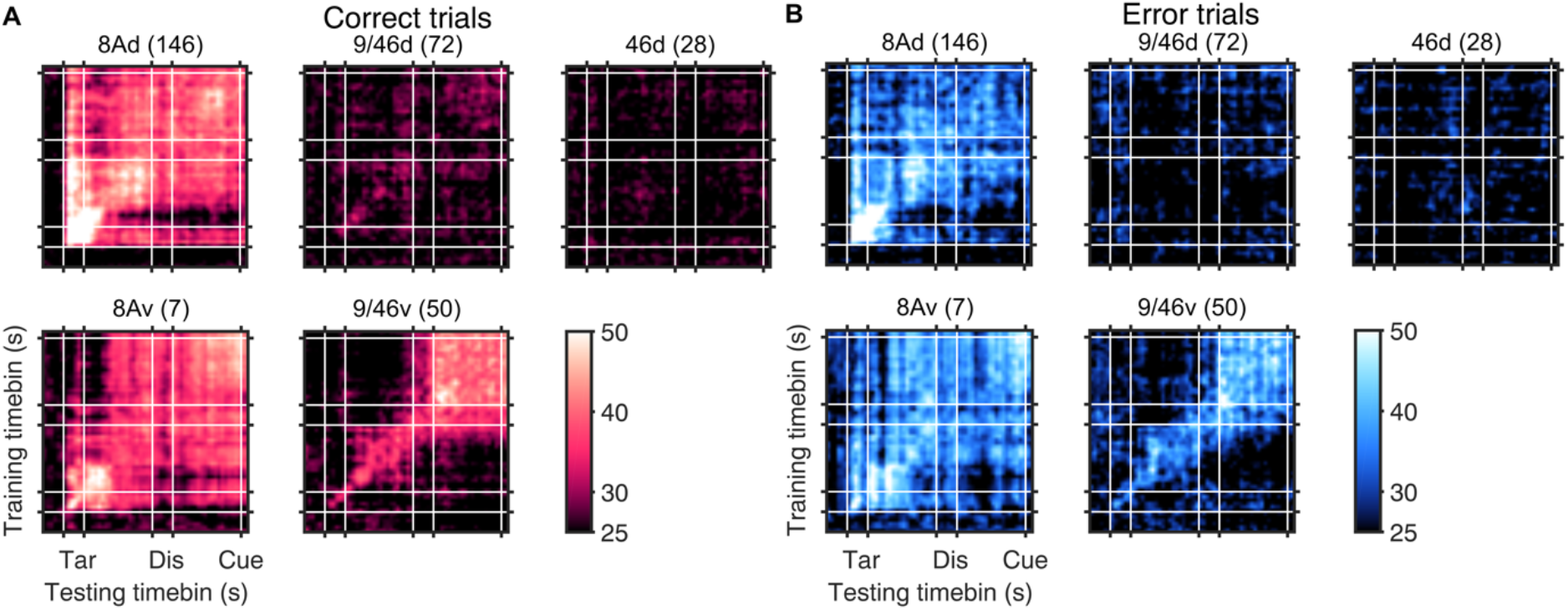
Decoding of 4 target locations for Monkey 1 only, with all neurons included per region. Since only Monkey 1 contributes neurons to area 46d, the lack of information in this area may be due to a lack of target information across regions in this monkey. To assess this possibility, we tested whether target information could be decoded from Monkey 1. The figure shows that all regions contain target information, except for areas 9/46d and 46d. The value in brackets beside the region names is the number of neurons used for decoding, which is also the total neurons recorded in Monkey 1 in that region. **A**, Decoding target location in correct trials, **B**, Decoding in error trials.

**Figure 8 – figure supplement 3.**
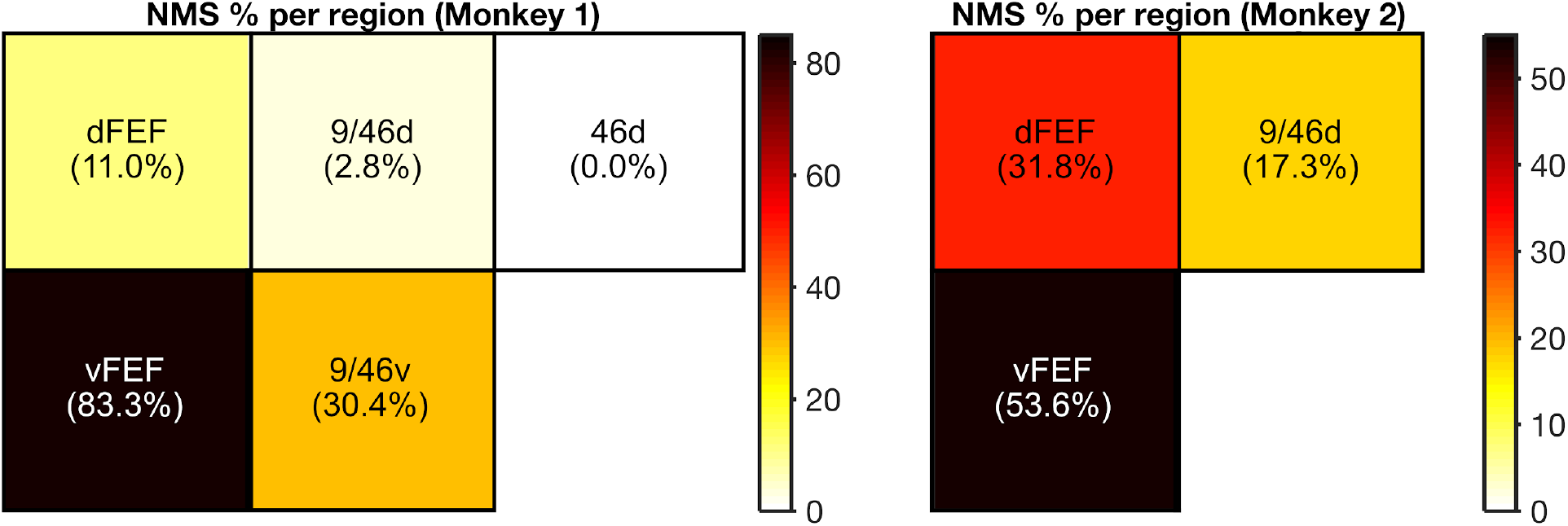
Proportion of nonlinear mixed selective cell proportions per region for each monkey. Only regions with recorded cells are shown for each animal.

**Figure 10 – figure supplement 1.**
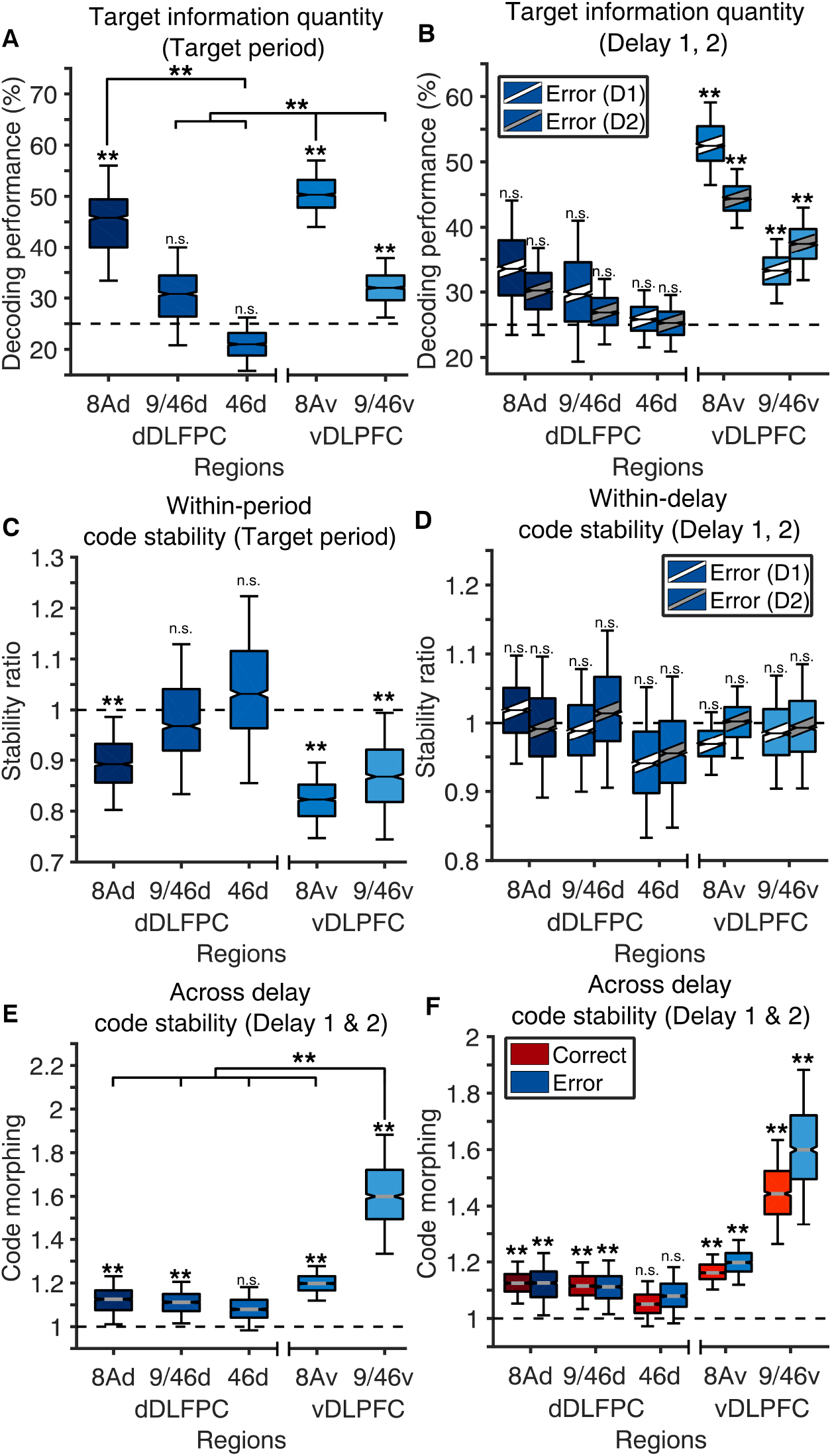
Regional differences in Delay 1 and 2 target information quantity and stability in error trials. A. , Information quantity in target period for the above regions, **B**, Information quantity in delays 1 and 2, **C**, Information stability in target period, **D**, Information stability in delays 1 and 2, **E**, Code stability across delays 1 and 2, **F**, Code stability across both delays 1 and 2 combined, in correct and error trials.

**Figure 10 – figure supplement 2.**
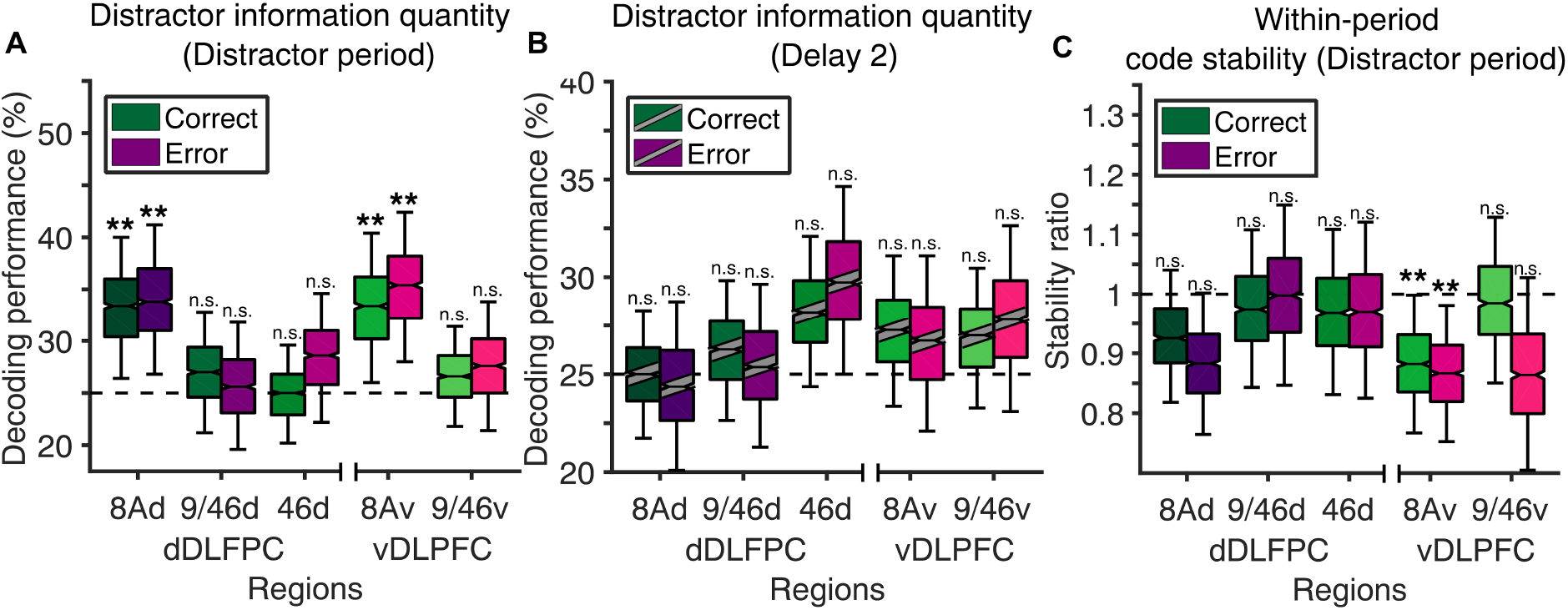
Differences in distractor information quantity and stability is not behaviorally relevant. **A**, Information quantity in the distractor presentation period for the above regions, **B**, Information quantity in Delay 2, **C** Information stability in Delay 2.

## Notes

### Competing Interest Statement

The authors have declared no competing interest.

